# Spotiphy enables single-cell spatial whole transcriptomics across the entire section

**DOI:** 10.1101/2024.11.11.623040

**Authors:** Jiyuan Yang, Ziqian Zheng, Yun Jiao, Kaiwen Yu, Sheetal Bhatara, Xu Yang, Sivaraman Natarajan, Jiahui Zhang, John Easton, Koon-Kiu Yan, Junmin Peng, Kaibo Liu, Jiyang Yu

**Affiliations:** Department of Computational Biology, St. Jude Children’s Research Hospital, Memphis, TN, 38105, USA; Department of Industrial & Systems Engineering, University of Wisconsin-Madison, Madison, WI, 53706, USA; Department of Structural Biology, St. Jude Children’s Research Hospital, Memphis, TN, 38105, USA; Center of Proteomics and Metabolomics, St. Jude Children’s Research Hospital, Memphis, TN, 38105, USA

## Abstract

Spatial transcriptomics (ST) has advanced our understanding of tissue regionalization by enabling the visualization of gene expression within whole tissue sections^1^, but the approach remains dogged by the challenge of achieving single-cell resolution without sacrificing whole genome coverage^2,3^. Here we present Spotiphy (<u>Spot</u> imager with <u>p</u>seudo single-cell resolution <u>h</u>istolog<u>y</u>), a novel computational toolkit that transforms sequencing-based ST data into single-cell-resolved whole-transcriptome images. In evaluations with Alzheimer’s disease (AD) and normal mouse brains, Spotiphy delivers the most precise cellular compositions. For the first time, Spotiphy reveals novel astrocyte regional specification in mouse brains. It distinguishes sub-populations of DAM (Disease-Associated Microglia) located in different AD mouse brain regions. Spotiphy also identifies multiple spatial domains as well as changes in the patterns of tumor-tumor microenvironment interactions using human breast ST data. Spotiphy enables visualization of cell localization and gene expression in tissue sections, offering key insights into the function of complex biological systems.

## Main

The location of a cell within its microenvironment is a critical determinant of its function and interactions with neighboring cells^4–6^. Cells are endowed with unique characteristics and functions that are finely tuned to meet the specific demands of their respective environments through a process called cellular regional specification^7^. Analyzing local cell behavior involves identifying spatial domains: regions characterized by cellular composition and their transcriptomic profiles. Cells’ reactions to external signals are influenced by their spatial domain, affecting tissue development, homeostasis, and disease progression^8^. However, current ST platforms^2,3,9,10^ can’t provide both the localization and transcriptomic data needed for this analysis. ST technology can generally be grouped into two categories: sequencing-based approaches (e.g., Visium^11^, DBiT-Seq^12^, Slide-Seq^13,14^, Slide-tags^15^, Stereo-seq^16^) and image-based approaches (e.g., SeqFISH^17,18^, smFISH^19^, RNAScope^20^, MERFISH^21–24^, Xenium^25^, and CosMx^26^). Pre-defining a capture area, or “spot”, allows sequencing-based approaches to generate unbiased genome-wide transcriptomic profiles while minimizing lateral leaking contamination. Yet limiting the reads to predetermined spots comes with its own major drawbacks: loss of capture in the area between the spots (non-capture areas) and low resolution (as each spot consists of multiple, often heterogenous, cells). Image-based ST approaches using high-resolution fluorescence images achieve single-cell resolution but are limited to pre-selected gene panels consisting of 300-1000 targets. They usually capture limited counts per gene, making them better for validation than data-driven discovery. Initial attempts focused on enhancing the resolution of sequencing-based ST data, with various deconvolution methods for spot-level ST data already established. These methods integrate scRNA-seq and ST data to parse the cellular composition of individual transcriptomic spots^27–41^. Yet, none of the current methods are able to achieve single-cell resolution while maintaining the spatially varying gene expression present in the original sequencing-based ST data.

Here we present Spotiphy, a toolkit that resolves the gene coverage-resolution tradeoff. Spotiphy generates inferred single-cell RNA expression profiles (iscRNA data) of *all* cells (located in *both capture and non-capture areas*) to achieve spatially resolved whole-slide transcriptomic profiling, providing substantial benefits for downstream analyses and opportunities for biological insight. Spotiphy-generated iscRNA data revealed regional specification of astrocytes and microglia in normal and AD mouse brains, a level of novel detail not detectable by existing scRNA-seq and ST technologies. Spotiphy identifies changes in the patterns of tumor-tumor microenvironment (TME) interactions across multiple spatial domains. It delivers single-cell-resolved whole-transcriptome images of the entire section that significantly expands the information intensity of ST data, providing a more comprehensive and detailed understanding of spatially resolved cellular states and regulatory mechanisms.

## Results

### Spotiphy achieves single-cell spatial whole transcriptomics via generative modeling

Spotiphy delivers single-cell spatial whole transcriptome profiling via generative modeling of sequencing-based ST, scRNA-seq, and histological imaging data (Fig. 1a, Methods). Specifically, Spotiphy selects the most informative genes for each cell type to generate a signature reference from scRNA-seq data. It presets five customizable hyperparameters to ensure scRNA-seq reference accuracy and robustness (Supplementary Fig. 1). Spotiphy also determines the locations of the nuclei by segmenting the high-resolution histological images. Spotiphy then integrates the above information into a probabilistic model that considers the distribution of gene contributions from each cell type. This unique feature enables the simultaneous deconvolution and decomposition of ST data, thereby generating both cell-type proportion and iscRNA data. It then imputes these data for cells located in non-capture areas through the Gaussian processes^42^. Consequently, Spotiphy generates single-cell-resolved images equivalent to the output of image-based approaches, with whole transcriptomic profiling across the entire sections.

**Fig. 1.**
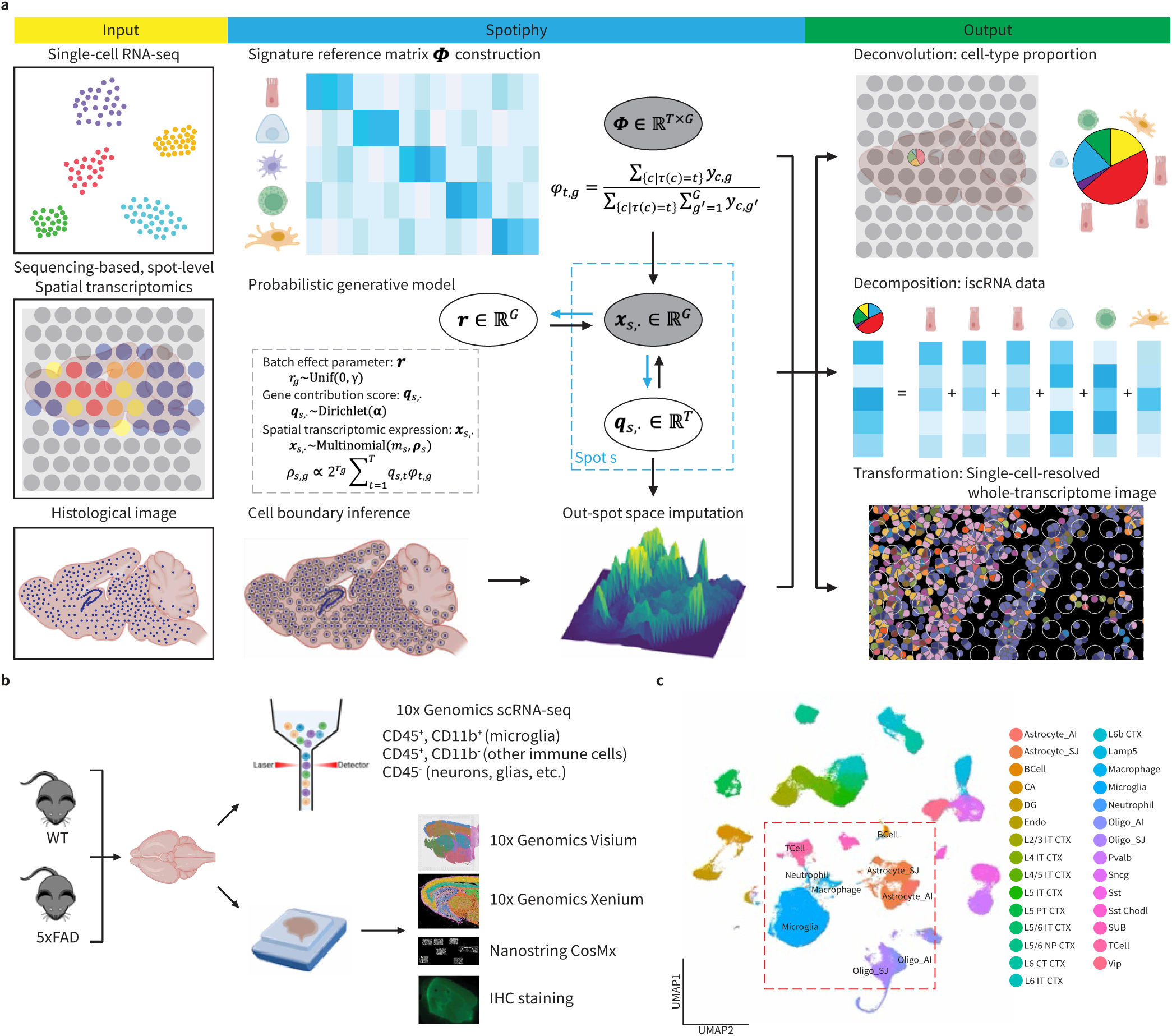
Schematic overview of Spotiphy and the matched datasets used for validation. **a** Workflow summary for Spotiphy. Spotiphy requires three types of input data: scRNA-seq data, spot-level sequencing-based ST data, and high-resolution histological images. The workflow consists of five major steps: (i) Reference construction: scRNA-seq data is used to select top marker genes and generate a signature reference for all cell types; (ii) Segmentation: the histological image is used to count the number of nuclei in each spot; (iii) Generative modeling: the ST data is used to generate cell-type proportions and iscRNA data; (iv) Imputation: Gaussian processes are used to impute the cell-type proportions and iscRNA data for cells located outside of the transcriptomic spots; (v) Transformation: the results of steps iii-iv are merged to produce a pseudo single-cell-resolved whole-transcriptome image. **b** The process of generating matched datasets for WT and AD mouse brain tissues. Some icons are from BioRender. **c** UMAP projection of the scRNA-seq reference data for mouse brain. 22 cell types from the Allen Institute atlas and 7 cell types from our original data (red box) are included. Astrocytes and oligodendrocytes from both datasets are included for batch correction. Astrocyte_AI: astrocytes from the Allen Institute atlas, Astrocyte_SJ: astrocytes from original data; Oligo_AI: oligodendrocytes from the Allen Institute atlas; Oligo_SJ: oligodendrocytes from the original data; CA: cornu ammonis; DG: dentate gyrus; Endo: endothelial.

### Matched mouse brain datasets for biological validity

To evaluate Spotiphy’s performance and the biological validity of its results, we enriched rare immune cells and obtained scRNA-seq profiles for 27,836 CD45^+^ and CD11b^+^ cells (microglia, macrophages, and neutrophils), 6,085 CD45^+^ and CD11b^-^ cells (T cells, B cells), and 29,563 CD45^-^ and CD11b^-^ cells (neurons, glial cells, etc.) from an AD mouse model^43^ and wild-type control (Fig. 1b, Methods). By supplementing our data with the Allen Institute’s atlas^44^, we assembled a comprehensive mouse brain cell reference with twenty-seven cell types including neurons, glia, and immune cells (Fig. 1c, Supplementary Fig. 2a-b). We then generated datasets of various ST techniques including IHC, Visium, Xenium, and CosMx. Serial sectioning of the same sample produced nearly identical slices, allowing direct comparison between different ST approaches once the histological images from these platforms were aligned (Supplementary Fig. 2c-e). The matched datasets are a valuable resource for the spatial omics community, particularly for the development and evaluation of ST algorithms.

### Spotiphy provides exceeding accuracy in deconvolution for rare cell-type proportion

To evaluate the accuracy of Spotiphy’s cellular deconvolution, Xenium data generated from the AD sample was used as the ground truth and the heatmaps of cell-type proportions across section were generated by Spotiphy and thirteen other benchmarking methods (Fig. 2a, Extended Data Fig. 1a, Supplementary Fig. 3-5, Methods). For well-organized excitatory glutamatergic neurons, Spotiphy’s delineation of the cortex layers was closer to that of Xenium and had clearer boundaries than those of the other methods. Spotiphy’s distribution patterns for unevenly distributed inhibitory GABAergic interneurons were likewise closer to the ground truth. Because the Xenium panel did not include markers for immune cells, we used the *in-situ* hybridization (ISH) results from the Allen Institute as our ground truth for neutrophils, T and B cells^45^. Whereas most other methods being tested failed to predict the distribution of neutrophils and instead producing random results, Spotiphy identified a specific enrichment of neutrophils around the ventricle that is in line with the ISH results for neutrophil markers, including *Ltgb2* (Fig. 2a, Supplementary Fig. 4-6). RCTD appeared comparable to Spotiphy, yet it incorrectly classified some neutrophils as macrophages, leading to an apparent increase of macrophages around the ventricle that did not align with the Xenium data (Supplementary Fig. 4e). Similar misidentifications among microglia, macrophages, and neutrophils were observed in CIBERSORTx’s results (Supplementary Fig. 5e). No clear ISH signal was observed for *Cd3e* (T cell marker) or *Cd19* (B cell marker), consistent with the fact that T and B cells are rarely found in mouse brain. Due to the low numbers of immune cells, and varying cell counts per spot, using proportions for rare cells can lead to inaccuracies. Thus, we translated these proportions into absolute cell counts (multiplying by the total cells per spot segment) for comparison (Supplementary Fig. 7). For B cells, most methods performed well except CytoSPACE, StereoScope, iStar, and Cell2location. Spotiphy exhibited a low false-positive detection of T cells in the striatum (upper right region), its prediction for macrophage and microglia aligned well with the ground truth. Multiple metrics were used to benchmark Spotiphy’s cellular deconvolution against other methods (Fig. 2b-d, Extended Data Fig. 1b-e, Supplementary Fig. 3). Spotiphy consistently produced the highest overall Pearson’s correlation coefficient with the ground truth. Spotiphy’s cell-type proportions for each transcriptomic spot also aligned more closely with the ground truth, as demonstrated by the correlation, the fraction of cells correctly mapped, and the cosine similarity. Furthermore, Spotiphy’s results also produced significantly lower values for the absolute error, mean square error, and Jensen–Shannon divergence (JSD).

**Fig. 2.**
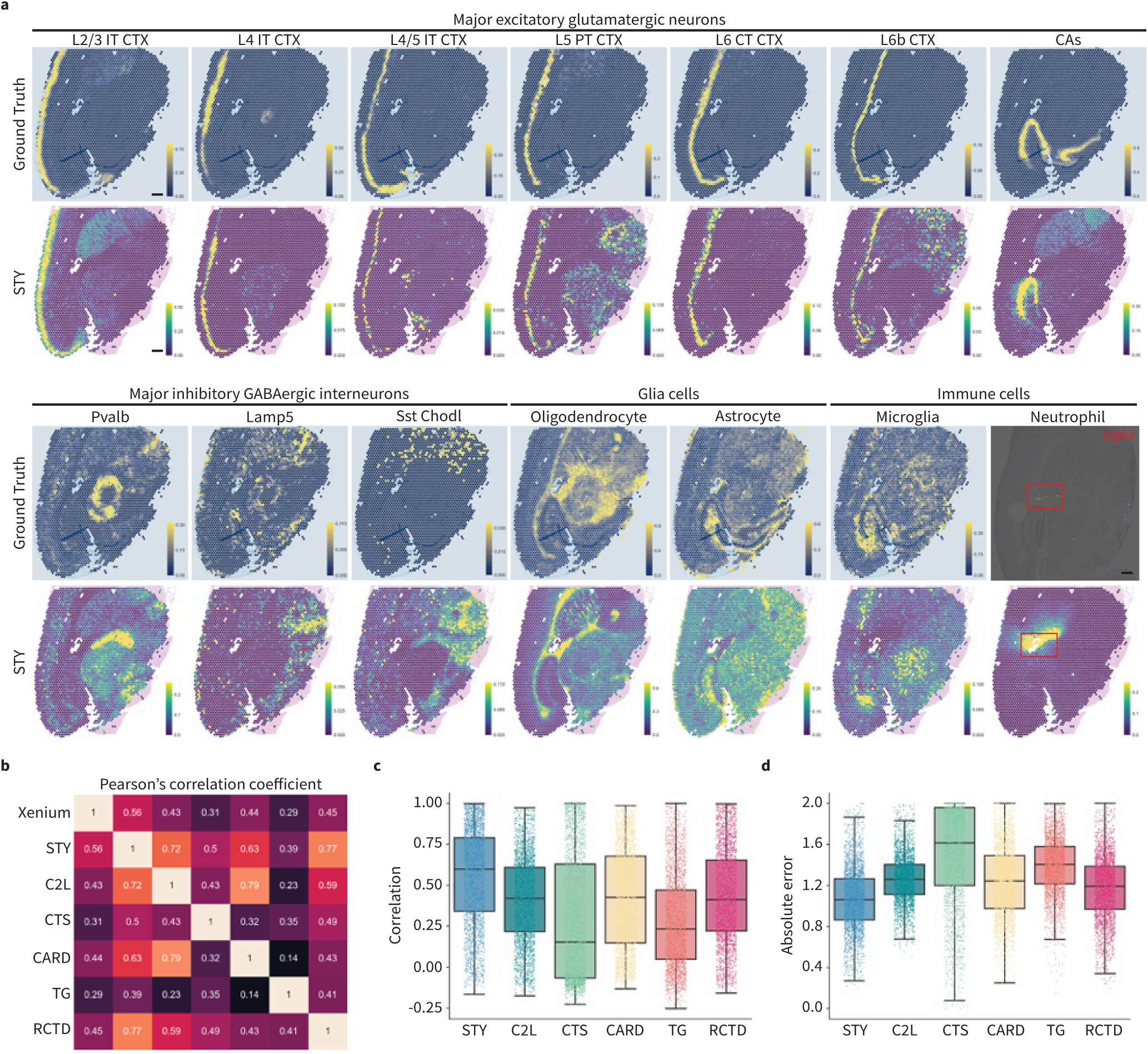
Benchmarking Spotiphy’s cellular deconvolution using matched Xenium data. **a,** Heatmaps depicting the proportion of selected cell types (as determined by Spotiphy and Xenium, the ground truth) across the histological image of AD sample. Red box in results of Neutrophil represent the ventricle region. Scale bar: 500 μm. **b**, Pearson correlation coefficient heatmap of cell-type proportions generated by Xenium, Spotiphy, and other methods selected for benchmarking. STY: Spotiphy, C2L: Cell2location, CTS: CytoSPACE, TG: Tangram. Box plots for the correlation (**c**, higher is better) and absolute error (**d**, lower is better) of the cell-type proportions for each transcriptomic spot generated by each method.

While the matched Xenium data was useful for evaluating the biological validity, it was not an ideal choice of ground truth for benchmarking the method’s performance. This is because (i) Xenium’s pre-designed gene panel has limited plexity, and (ii) the data was generated from serial section instead of identical one. We therefore constructed a series of simulated Visium datasets from scRNA-seq data with various noise levels to obtain a comprehensive benchmarking analysis, and Spotiphy continued to demonstrate the best performance (Extended Data Fig. 2a-e, Supplementary Fig. 8-10, Methods). Additionally, eight paired “scRNA-Visium” datasets from various human tissues^41^ were used to further evaluate the deconvolution performance of Spotiphy against the other thirteen methods. The quantitative metrics further confirmed that Spotiphy exhibited the lowest error and highest correlation to the ground truth (Supplementary Fig. 11). Moreover, Spotiphy’s computational time was substantially lower than that of the other methods (Extended Data Fig. 2f). In summary, with a total of thirteen methods included for deconvolution benchmarking, our evaluations demonstrate Spotiphy’s superiority across different tissue types.

### Spotiphy captures novel astrocyte regional specification in mouse brain tissue

Besides surpassing its peers in deconvolution, Spotify is unique in its ability to decompose the transcriptomic profiles of spots into single-cell level (iscRNA data, Extended Data Fig. 3a, Methods). When we applied unsupervised clustering^46^ to the iscRNA data of 33,819 cells generated from the Visium data of mouse brains, individual astrocytes did not cluster together; rather, they formed multiple sub-clusters, indicating intra-heterogeneity not apparent in the single-cell reference (Fig. 1c, Extended Data Fig. 3b, Supplementary Fig. 12, Fig. 3a). Similar phenotypes were observed in oligodendrocytes and multiple types of neurons. We subsequently color-coded the astrocytes based on their sub-cluster and superimposed them on the histological images according to their transcriptomic spot-of-origin (Fig. 3b). Surprisingly, the distinct spatial distribution of these sub-clusters corresponded perfectly with the classical histological regions of the mouse brain, in which the sagittal section is divided into six major topographic regions: cerebral cortex (CTX), hippocampus (HPF), fiber tracts (FT), thalamus (TH), hypothalamus (HY), and striatum (STR)^47,48^ (Extended Data Fig. 3f). Notably, the spatial distribution of astrocyte sub-clusters was nearly identical in the WT and AD samples; the proportions of each astrocyte subtype likewise did not differ substantially between the two samples (Fig. 3b, Extended Data Fig. 3e).

**Fig. 3.**
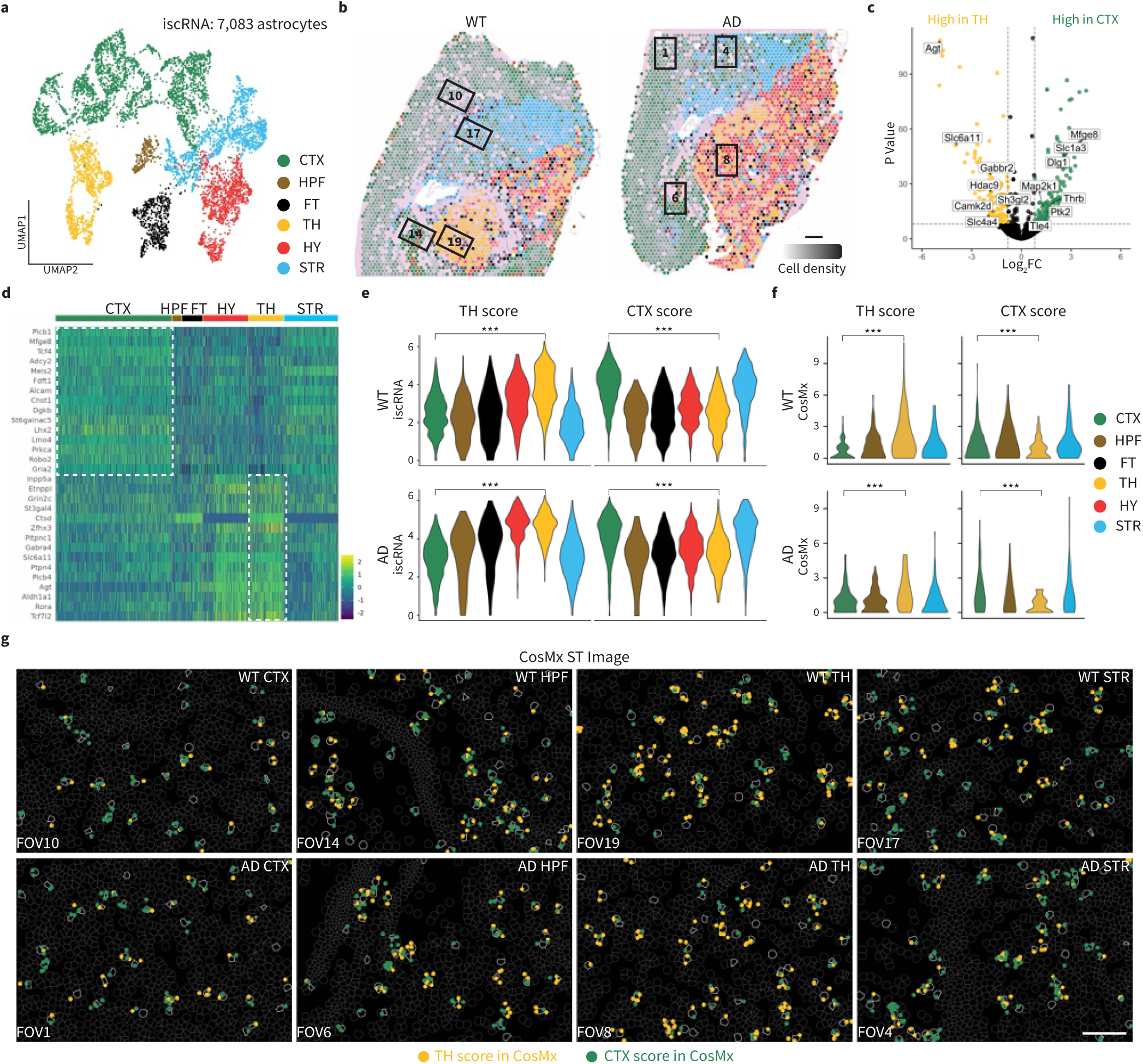
Spotiphy captures astrocyte regional specification in mouse brain tissue. **a,** UMAP projection of 7,083 astrocytes extracted from iscRNA data produced by applying Spotiphy to mouse brain Visium data. Clusters are labeled according to their corresponding topographic regions. CTX: cerebral cortex; HPF: hippocampus; FT: fiber tracts; TH: thalamus; HY: hypothalamus; and STR: striatum. **b**, Transcriptomic spots from the Visium data. Spots are color-coded according to their constituent astrocytes’ subcluster annotated in **a**. The opacity of the spots is representative of cell density. Spots not containing astrocytes have been removed for clarity. Black boxes represent the corresponding FOVs of the CosMx data. FOV1: AD CT; FOV6: AD HPF; FOV8: AD TH; FOV4: AD STR; FOV10: WT CTX; FOV14: WT HPF; FOV19: WT TH; and FOV17: WT STR. Scale bar: 500 μm. **c**, Volcano plot of differentially expressed genes (DEGs) among astrocytes in the CTX vs. TH. DEGs were determined using the iscRNA data. The genes highlighted are available in the CosMx panel and have been used to generate the signature scores in **e-g**. **d**, Heatmap of the expression of the top 15 DEGs among astrocytes in the CTX region and the TH region, respectively. **e-f**, Violin plots of the CTX and TH scores for each astrocyte sub-cluster identified from the iscRNA (**e**) and CosMx (**f**) data. Two-sided t-tests were conducted to determine the statistical significance of the CTX and TH scores of the astrocytes in each sub-cluster. ***, p < 0.01. **g**, Visualization of signature genes in selected FOVs of the CosMx data. Yellow and green dots represent the signature genes used to calculate the TH and CTX scores, respectively. Cell boundaries are depicted with gray lines; astrocytes are outlined in white. Scale bar: 100 μm.

Further downstream analysis of the iscRNA data generated by Spotiphy revealed regional specification of astrocytes. To explore potential functional differences in the astrocytes of the telencephalon and diencephalon, we selected a region from each (CTX and TH, respectively)^48^ for comparison. Using NetBID2^49^, we identified differentially expressed genes (DEGs) between CTX and TH astrocytes (Fig. 3c-d) and then used the top DEGs to define signature scores for each region (TH score and CTX score). As expected, each astrocyte scored high for its region-of-origin and low for the comparison region in both WT and AD mouse samples (Fig. 3e). To validate the accuracy of the iscRNA data and downstream observations, we used the CosMx data for the CTX and TH regions in the WT and AD mouse samples as the ground truth. In that data as well, astrocytes in the CTX region had a higher CTX score, and astrocytes in the TH region had a higher TH score (Fig. 3f-g), supporting the validity of the iscRNA data generated by Spotiphy. Differences in the actual transcripts (as measured by CosMx) of the astrocytes from these regions further supported this astrocyte regional specification. Gene set enrichment analysis (GSEA) demonstrated enrichment of CTX astrocytes in pathways associated with neuronal differentiation and cell fate specification; TH astrocytes, meanwhile, were found have greater expression of genes related to neuronal synaptic plasticity. These data suggest distinct biological roles for astrocytes in different regions of the mouse brain (Extended Data Fig. 3g).

A recent study identified two distinct astrocyte clusters in mouse brains using snRNA-seq profiles of astrocytes enriched in different regions^28^. These clusters aligned with the telencephalon (Cluster I) and diencephalon (Cluster II), indicating regional expression differences in astrocytes (Extended Data Fig. 4a). This data is consistent with the astrocyte regional specification that we observed in the iscRNA data generated by Spotiphy. When we applied our signature scores to this data, astrocytes in Cluster I had a high CTX score and low TH score, whereas those in Cluster II had a low CTX score and a high TH score, supporting the biological validity of our iscRNA data (Extended Data Fig. 4b-c).

Furthermore, we generated iscRNA data from the simulated Visium datasets discussed above. Unsupervised clustering of the simulated iscRNA data showed that astrocytes, oligodendrocytes, and neurons were clustered together and did not form sub-clusters as they did in the real iscRNA data (Extended Data Fig. 4d-e). Nearly all astrocytes grouped into one single cluster, except for two minor subclusters, which exhibited random distribution and no clear regional patterns (Extended Data Fig. 4f-g). As the simulated Visium datasets were generated using scRNA-seq data, the lack of spatial variable genes and identifiable regional specification were anticipated. This result suggest that Spotiphy will not introduce any "variations" stemming from its modeling process, and therefore the regional specification observed in the real iscRNA data reflects true biological phenomena rather than merely an artifact of Spotiphy.

### Spotiphy reveals novel DAM regional specifications in AD mouse brain tissue

To confirm that the iscRNA data generated by Spotiphy for rare cell populations is likewise biologically meaningful, we analyzed the microglial populations, which formed five sub-clusters corresponding to the histological regions of the mouse brain, with a single cluster (C&H) encompassing both the CTX and HPF regions (Fig. 4a-b, Extended Data Fig. 5a). Both the WT and AD brain tissue showed similar distribution patterns of microglia in the FT, HY, and STR regions; however, microglia localizing in the C&H and TH regions were found to originate mainly from the AD sample (Extended Data Fig. 5b). To characterize these AD-specific microglial populations, we used NetBID2 to identify the top DEGs and found significant overlap between those genes and disease-associated microglia (DAM) markers^50–55^, suggesting that the microglia in these regions of the AD brain may in fact be DAM (Extended Data Fig. 5c). We then utilized commonly used microglia and DAM markers to define signature scores and applied them to the iscRNA data for the microglia from both samples (Extended Data Fig. 5d). As expected, the microglia scores for each region did not vary significantly between the WT and AD samples. The DAM score, however, was significantly higher in the AD brain, especially in the C&H and TH sub-clusters (Fig. 4c). These observations were validated by the CosMx data, in which the signal intensity of DAM markers was greater for microglia located in the C&H and TH regions of the AD brain (Fig. 4d-e). Furthermore, GSEA analysis found a strong enrichment of immune response pathways in AD-specific microglia, consistent with previous scRNA-seq studies of DAM populations^50–55^ (Extended Data Fig. 5e). Given the strong co-localization of DAMs and beta-amyloid noted in previous studies^52,54,56,57^, IHC staining was performed on both samples (Extended Data Fig. 5f, Methods). Beta-amyloid signals were substantially higher in the C&H, FT, and TH regions of the AD brain, with the remaining regions showing no significant difference from regions without microglia (labeled as n/a, Extended Data Fig. 5g). Together, these results support that the AD-specific microglia in the C&H and TH regions were indeed the DAM population.

**Fig. 4.**
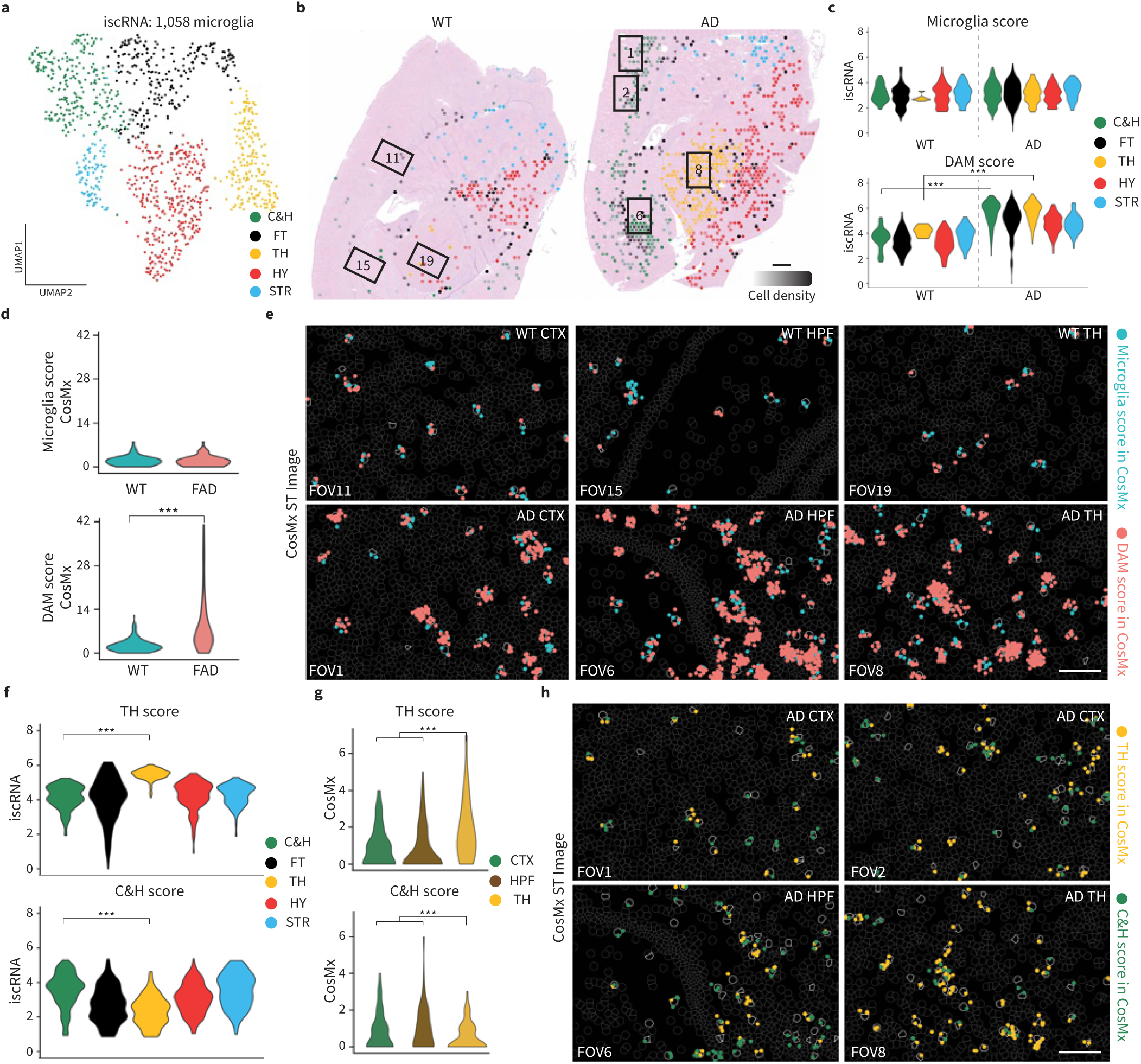
Spotiphy reveals microglia regional specification in AD mouse brain tissue. **a,** UMAP projection of 1,058 microglia extracted from iscRNA data generated by Spotiphy to mouse brain Visium data. Clusters are labeled according to their corresponding topographic regions. C&H: cerebral cortex and hippocampus. **b**, Transcriptomic spots from the Visium data. Spots are color-coded according to their constituent microglia’s sub-cluster, annotated in **a**. The opacity of the spots is representative of cell density. Spots without microglia have been removed for clarity. Black boxes represent the corresponding FOVs of the CosMx data. FOV1: AD CTX; 2: AD CTX; FOV6: AD HPF; FOV8: AD TH; FOV11: WT CTX; FOV15: WT HPF; FOV19: WT TH. Scale bar: 500 μm. **c**-**d**, Violin plots of the microglia and DAM scores for each microglia sub-clusters identified from the iscRNA (**c**) and CosMx (**d**) data. **e**, Visualization of signature genes in selected FOVs of the CosMx data. Blue and red dots represent the signature genes used to calculate the microglia and DAM scores, respectively. **f**-**g**, Violin plots of the C&H and TH scores for the microglia sub-clusters present in the FAD sample’s iscRNA (**f**) and CosMx (**g**) data. **h**, Visualization of signature genes in selected FOVs of the CosMx data. Yellow and green dots represent the signature genes used to calculate the TH and C&H scores, respectively. Cell boundaries are depicted with gray lines; microglia are outlined in white. Scale bar in **e, h**: 100 μm. Two-sided t-tests are conducted in **c-d** and **f-g**. ***, p < 0.01.

Unsupervised clustering of the iscRNA data for the AD sample identified C&H and TH sub-clusters of DAMs, indicating DAM regional specification (Extended Data Fig. 6a). This regional specification was supported by a further validation that used CosMx data as ground truth. NetBID2 analysis identified DEGs for the C&H DAM and TH DAM subclusters (Extended Data Fig. 6b-c); these were then used to define signature C&H and TH scores that were applied to both the CosMx and iscRNA data. As expected, the iscRNA and CosMx data for the C&H DAM had higher C&H scores, and both data for the TH DAM had higher TH scores (Fig. 4f-h). GSEA analysis showed that C&H DAM exhibit upregulation of genes related to the immune response, indicating greater immune activation in the CTX and HPF regions of the AD sample (Extended Data Fig. 6d). This finding was consistent with the greater relative accumulation of beta-amyloid that we observed in the CTX and HPF regions of the AD sample^58,59^. The microglia in the simulated iscRNA data were clustered together and had a seemingly random distribution, confirming that the results were authentic and not artifact-related (Extended Data Fig. 6e-f).

### Spotiphy charts spatial domains of tumor and TME in breast cancer

To evaluate Spotiphy’s ability to characterize tumors and the tumor microenvironment (TME), publicly available scRNA-seq and Visium datasets^60,61^ for breast tissue samples (BCCID4535, BC1160920F, BCCID44971) were applied to Spotiphy and produced iscRNA data for a total of 23,200 cells (Supplementary Fig. 13, Methods). Unsupervised clustering analysis of this iscRNA data revealed multiple sub-clusters for luminal hormone responsive (LumHR) cells and luminal secretory (LumSec) cells (Fig. 5a-b). To test if the iscRNA data could distinguish tumors from normal cells, we applied inferCNV analysis^62–66^ to both Visium and iscRNA data (Fig. 5c, Supplementary Fig. 14). Although the spot-level Visium data can reveal significant copy number variations (CNVs) in both tumor samples, obvious intra-heterogeneity in the CNV patterns was also observed in BC1160920F. Due to resolution limits, we cannot determine which cells contribute to this variation. In contrast, the inferCNV results of iscRNA data showed that LumHR cells from BCCID4535 and LumSec cells from BC1160920F possessed more copy number variations (CNVs), indicating their greater likelihood of being tumor cells. Both LumHR and LumSec cells from BCCID44971 exhibited fewer CNVs, indicating they are normal cells. This observation was consistent with the original study’s conclusion that BCCID44971 was normal tissue^61^. We subsequently validated the tumor/non-tumor identities of the cells (as determined by the iscRNA data) using Resolve smFISH data from tumor (P35-S1) and normal (P69-S3) breast tissue samples^60^. We selected the DEGs for tumor (BC1160920F) and normal (BCCID44971) LumSec cells and then used them to define T-Sec (tumor LumSec) and N-Sec (normal LumSec) scores, which we subsequently applied to the iscRNA and smFISH datasets (Extended Data Fig. 7c-d). Indeed, genes that had elevated expression in tumor LumSec cells also produced greater signal intensity in the smFISH data for the tumor sample (P35-S1); the same was observed for the N-Sec genes (Fig. 5f-h). Similarly, the top DEGs for the tumor (BCCID4535) LumHR cells were also found to have elevated expression and greater signal intensity in both the iscRNA data and the smFISH data for the tumor sample (P35-S1), respectively (Extended Data Fig. 7e-i). Furthermore, GSEA analysis found elevated immune activation features in the cells originating from the tumor samples (BC1160920F and BCCID4535) (Extended Data Fig. 7j-k), supporting Spotiphy’s accuracy in differentiating between tumor cells and normal cells.

**Fig. 5.**
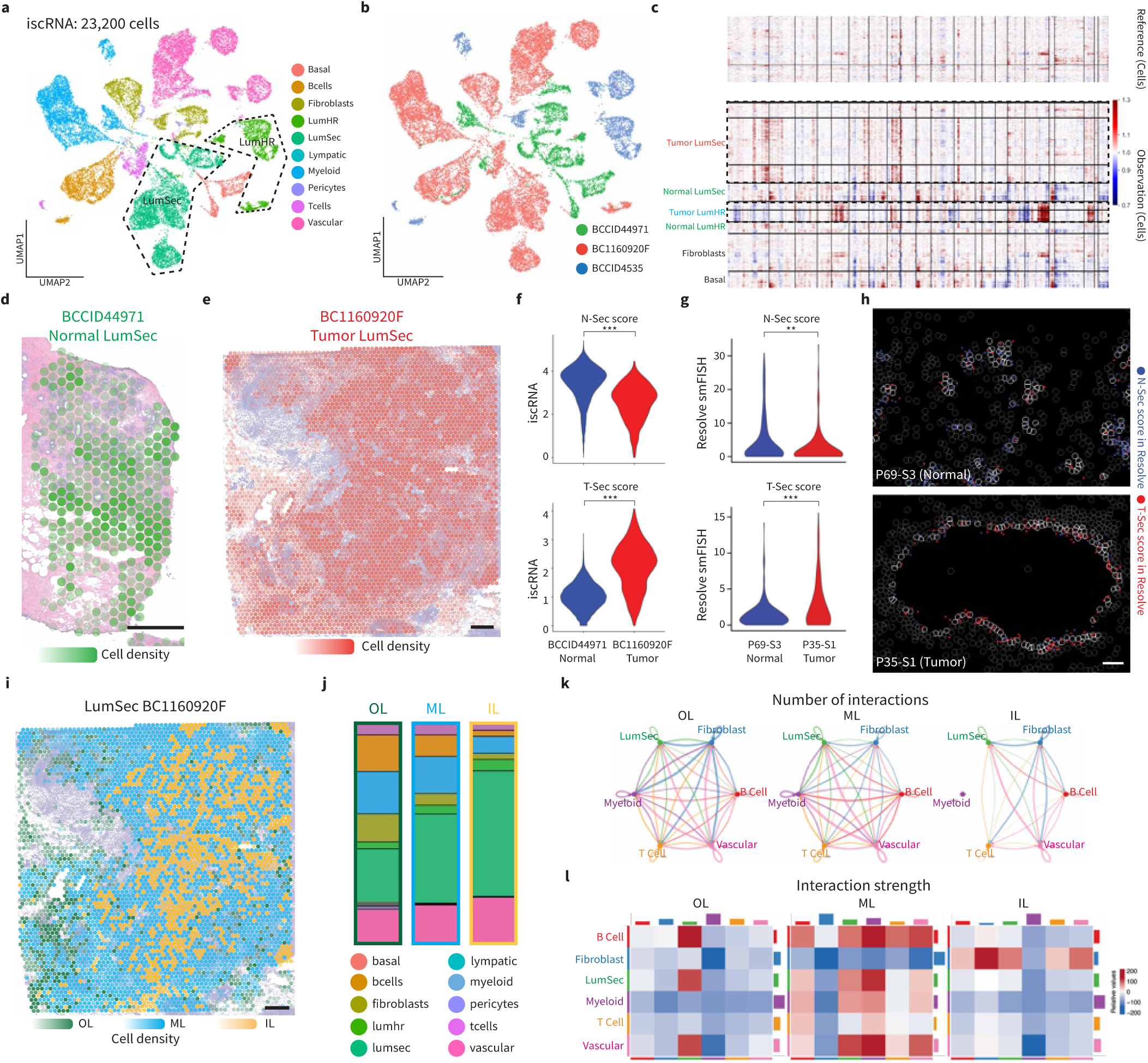
Spotiphy reveals tumor-TME changes across spatial domains in human breast tissue. **a-b,** UMAP projection of 23,200 cells from iscRNA data produced by applying Spotiphy to human breast tissue Visium data. Cells are color-coded according to their cell type (**a**) or sample-of-origin (**b**). LumSec and LumHR cells are encircled by dashed black lines in **a**. **c**, The results of applying inferCNV to the iscRNA data in **a**. Tumor cells originating from BC1160920F (LumSec) and BCCID4535 (LumHR) are encircled by dashed black lines. **d-e**, Transcriptomic spots from the Visium data are color-coded according to their constituent LumSec cells’ sample-of-origin: BCCID4491 (d) and BC1160920F (e). The opacity of the spots is representative of cell density. Scale bars: 500 μm. **f,** Violin plots of the N-Sec and T-Sec scores for the LumSec cells identified in the iscRNA data of normal sample BCCID4491 and tumor sample BC1160920F. **g,** Violin plots of the N-Sec and T-Sec scores for the LumSec cells identified in the Resolve smFISH data of normal sample P69-S3 and tumor sample P35-S1. Two-sided t-tests were conducted. ***, p < 0.01, **, p < 0.05. **h**, Visualization of signature genes in selected regions of the Resolve smFISH data. Blue and red represent the signature genes used to calculate the N-Sec and T-Sec scores, respectively. Cell boundaries are depicted with gray lines; LumSec cells are outlined in white. Scale bars: 100 μm. **i,** Transcriptomic spots from the Visium data. Spots are color-coded according to their constituent LumSec cells’ spatial domains. Scale bars: 500 μm. **j**, Cell-type proportions of the three spatial domains based on the iscRNA data. **k-l**, Results of applying CellChat to the iscRNA data of the cells in the three LumSec spatial domains. **k**, Number of interactions among cell types across three spatial domains. **l**, Interaction strength among cell types across three spatial domains. OL: outer layer; ML: middle layer; and IL: inner layer.

Three LumSec tumor cell sub-clusters in BC1160920F exhibited unique localizations, distributed from center to periphery of the section (Extended Data Fig. 8a, Fig. 5i). We categorized spots into three “spatial domains” based on the sub-cluster of their LumSec cells, termed as outer layer (OL), middle layer (ML), and inner layer (IL) (Fig. 5j). We labeled cells by its spatial domain and divided the iscRNA data into three domain-specific subsets. Similar distribution patterns were observed when overlaid OL and IL spatial domains on the BCCID4535 (Extended Data Fig. 8a-c). Interestingly, the cell-type proportions varied across the spatial domains, with tumor cells (LumSec and LumHR cells) increasing and immune cells (T, B, and Myeloid) decreasing towards the center (Fig. 5j, Extended Data Fig. 8c).

To better understand these variations, we used CellChat^67,68^ to compare the tumor-TME communication patterns of these spatial domains. Dramatic differences were observed in the tumor-TME interactions for both BC1160920F and BCCID4535 (Fig. 5k-l, Extended Data Fig. 8d-e). Interestingly, the interaction strength (per CellChat) between tumor and immune cells didn’t always correspond to the cell-type proportions of the spatial domain. For LumSec cells in BC1160920F, the strongest immune reactions were detected in the ML domain, not the OL domain with the most immune cells (Fig. 5l). Interactions between "collagen family – SDC1/4" and "MDK – SDC1/4" are associated with cell mobility, were notably increased in the ML domain ^69,70^. Both pairs were upregulated in the ML domain of BC1160920F and the OL domain of BCCID4535 (Extended Fig. 8f-g), indicating that cell mobility may play a role in the infiltration of immune cells into tumors.

To emphasize the novelty and superiority of Spotiphy’s decomposition strategy, we performed a comprehensive decomposition benchmarking of Spotiphy against Tangram, SpatialScope, and iStar using simulated Visium datasets of mouse brain (Supplementary Fig. 15). Four methods delivered cell-type-level expression matrices for each spot. Spotiphy consistently achieved the highest correlation and the cosine similarity, and lowest absolute error, mean square error, and JSD compared to the ground truth. Spotiphy also delivered the highest Matthews correlation coefficient (MCC) for each spot, indicating the superiority of Spotiphy’s decomposition performance. In addition, Spotiphy requires significantly less computation time than its competitors. Considering that a typical Visium sample usually contains 3000 to 4000 spots, this is a major advantage, allowing Spotiphy to deliver reliable results in a shorter time frame.

Another strategy for decomposition benchmarking involves merging the iscRNA data into pseudo-Visium data according to the spot-origin of cells and then comparing it with the original inputs (Supplementary Fig. 16). Since Spotiphy’s decomposition strategy involves partitioning gene counts into different cells, merging its iscRNA data results in data that is almost identical to the input, achieving the highest overall Pearson’s correlation coefficient. iStar also delivers good performance with the second highest PCC. Tangram and SpatialScope, however, demonstrated less than ideal results. We then examined the spatial distribution of six spatial variable genes (SVGs) in the pseudo-Visium data. Spotiphy’s output accurately preserves the original distribution patterns. As SpatialScope can only decompose the marker genes from its early phase, it only provided the distribution of two genes. This limitation significantly restricts its application compared to Spotiphy. In contrast, though Tangram’s iscRNA data have the whole-genome coverage, it completely distorted two genes’ distribution and exhibited less overall accuracy.

Collectively, these results strongly illustrate Spotiphy’s advancements, demonstrating its novelty, accuracy, and effectiveness. It brings single-cell resolution to the whole-transcriptomic ST data, preserving the unique distribution patterns of SVGs and creating opportunities for new insights.

### Spotiphy generates pseudo single-cell whole-slide spatial images

For the first time, Spotiphy addresses another major drawback of sequencing-based ST approaches: the information loss for the space between the transcriptomic spots (non-capture areas). Visium^11^ and DBiT-Seq^12^ typically sequence about 50% of the entire section. In both mouse brain Visium datasets, only 34% of cells were within the spots, leading to profound information loss (Extended Data Fig. 9a). After analyzing the seamless image-based Xenium datasets of human lung cancer, colorectal cancer, and mouse brain, we observed that within small scales (< 100 μm), cellular proportion changes are minimal. This allows us to impute the information for non-capture areas based on the nearby regions (Supplementary Fig. 17-18). In fact, cells of the same type typically exhibit a gradually uniform distribution within a tissue section to maintain tissue function^71–73^, a pattern also reported using the MERFISH platform^74^. Most genes mirror the steady and consistent distribution, particularly those involved in maintaining basic cellular functions and tissue architecture^75,76^. Consequently, Spotiphy imputes cell-type proportions of non-capture areas using its outputs via Gaussian processes (Extended Data Fig. 9b-c).

We then validated the imputation’s accuracy using two fields of view (FOVs) located in CTX and HPF in matched Visium and Xenium data from AD brain. CytoSPACE^30^ and Tangram^32^ were used to benchmark Spotiphy’s performance (Fig. 6a, Extended Data Fig. 9d). Coloring by cell types, Spotiphy yields pseudo-single-cell-resolution images that are highly consistent with the ground truth (Fig. 6b, Extended Data Fig. 9e). All methods delivered reasonable cell-type proportions for the spots, with Spotiphy’s results being closest to the ground truth (Fig. 6c-d). Intriguingly, the inferred cell-type proportions for the non-capture areas from Spotiphy was closer to the ground truth than the Tangram-and CytoSPACE-generated data for the ***spots***. Furthermore, the merged cell-type proportions for spots and non-capture areas matched the accuracy of its spot-only results. These findings support imputation via Gaussian processes as an approach to reconstructing the whole section from sequencing-based ST spot grids. Spotiphy also provides kernel density smoothing to estimate the transcriptomic profiles of cells in the non-capture areas (Methods).

**Fig. 6.**
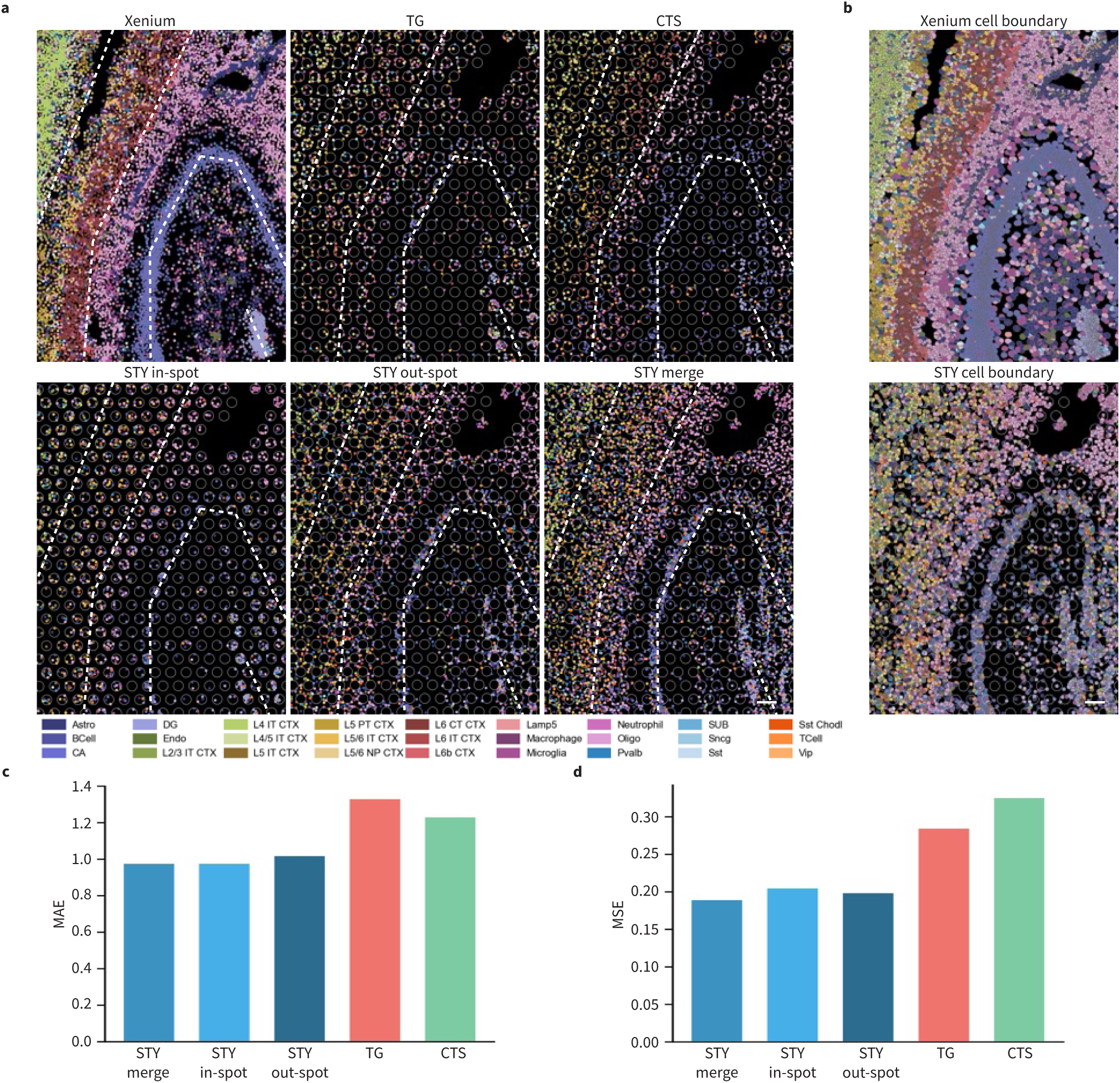
Spotiphy outputs pseudo single-cell-resolved whole-transcriptome images. **a-b,** Cell type annotation of individual nuclei and cells in selected FOVs at hippocampus. Dots in **a** represent all detectable nuclei color-coded for cell type. Grey circles represent the transcriptomic spots in the Visium data. Dashed white lines have been added to allow for easier comparison. Cells in **b** were color-coded for cell type. The cell boundaries depicted for the Xenium (upper panel) and Spotiphy (lower panel) data were inferred through those methods’ respective pipelines. Scale bars in **a-b**: 100 μm. **c-d**, The mean absolute error (MAE) and mean squared error (MSE) of the cell type imputation generated by Spotiphy (STY), Tangram (TG) and CytoSPACE (CTS) for each cell. Xenium data was used as ground truth. STY in-spot: image with only cells inside the spots; STY out-spot: image with only cells outside the spots; STY merge: image with all cells included.

### Spotiphy overcomes the limitations of *in situ* single-nuclei sequencing

Slide-tags is another sequencing-based platform recently introduced^15^. It labels individual nuclei in a whole section with unique barcodes, generating snRNA-seq data with high gene coverage along with spatial information. While Slide-tags represents an important innovation in the ST field, its practicality is limited by the method’s significant loss of nuclei (approximately 75%). Similar to how Spotiphy’s imputation function recovers data from non-capture areas in spot-based ST, treating each nucleus in Slide-tags data as a spot enables us to retrieve the lost snRNA-seq data. To pinpoint the “missing” nuclei that Slide-tags has not labeled, we aligned the H&E image of mouse hippocampus data (839 nuclei) from the original paper with the image of the same brain region in our AD brain, wherein 3,193 nuclei were detected (Extended Data Fig. 10a-c, Methods). Spotiphy provided iscRNA data for 3,193 nuclei in the AD brain using Slide-tags snRNA-seq data. Cells from both datasets mixed well by cell type after unsupervised clustering, indicating that Spotiphy’s results were accurate and free of batch effects (Extended Data Fig. 10d-e). The cell-type proportions obtained from Slide-tags snRNA-seq data and iscRNA data for the hippocampus were highly concordant, further supporting the accuracy of Spotiphy’s inference. The gene coverage of iscRNA data also closely matched the snRNA-seq data, showing no loss in coverage when applying Spotiphy to Slide-tags (Extended Data Fig. 10f-g).

Altogether, the experiments reported herein support the biological validity of iscRNA data generated by Spotiphy. These findings demonstrate that Spotiphy overcomes the key challenges of sequencing-based ST approaches, delivering accurate, biologically meaningful single-cell-resolved ST data compatible with computational packages for downstream analysis.

## Discussion

Here we introduced Spotiphy, an integrated method that marries the whole genome coverage of sequencing-based ST approaches with the single-cell resolution of image-based methods. Spotiphy offers a wider array of features (Supplementary Fig. 19a). Spotiphy’s superior inference of cell-type proportions^28–32^ largely owes to its improved marker gene selection and generative modeling approach. Rather than simply predict the cell-type proportions of transcriptomic spots, Spotiphy models the gene counts for each cell in each spot; in doing so, it not only significantly increases the information density, but also brings single-cell resolution to sequencing-based ST data. In contrast, Cell2location^28^ and RCTD^29^ employ probabilistic models without explicitly incorporating decomposed expression into their models. CytoSPACE^30^, CARD^31^, and Tangram^32^ approach deconvolution from an optimization perspective, attempting to identify the combination of cells in the scRNA-seq reference whose collective expression most closely resembles that of each spot in ST data. They face to two drawbacks: (i) Output accuracy heavily relied on the quality of reference data. If the reference lacks or only contains a limited number of the correct cell types, the algorithm would produce results significantly deviating from the ground truth. This explained their poor performance for immune cell populations. (ii) **No additional information** is gained from this mapping process, rendering them less powerful compared to Spotiphy. SpatialScope^27^ employs a strategy similar to RCTD for cell-type proportion deconvolution and enhances spot expression assembly with pseudo-cells that closely resemble the scRNA-seq reference, as generated by a deep-learning model. However, it confines the gene expression for each cell type to a predetermined range that aligns with the scRNA-seq reference, making it difficult to parse cellular regional specification. Spotiphy uses a Bayesian approach to dissect the expression of individual genes and assign them to specific cell types, producing iscRNA data that more accurately reflects expression fluctuations within transcriptomic spots and reveals cellular regional specifications. Capturing region-specific cell subtypes through conventional single-cell transcriptomics technologies is challenging for several reasons: i) The single-cell sampling may not cover enough regions of tissue, where ST data usually have full coverage of the whole tissue block. ii) The population of cells, particularly rare types of cells, is often too small to detect these intra-variations for one profiling. In comparison, Spotiphy can generate an average of about 10,000 cell expression profiles from a single tissue section (whereas one scRNA-seq sample typically has around 5,000 cells on average). iii) It is impossible to determine whether sub-clusters of certain cell types observed in scRNA-seq data are due to regional variations without the spatial information unique to ST data. For the first time, Spotiphy is able to recover the information loss for non-capture areas via Gaussian processes (Supplementary Fig. 20). The insights into the cellular organization, heterogeneity, and function within complex biological systems provided by Spotiphy will deepen our understanding of the molecular mechanisms underlying AD pathogenesis and other solid tumors.

Spotiphy’s results are reliant on the quality of input. Since the decomposition applies to every gene, the iscRNA data will retain the gene coverage of the ST data. Therefore, sequencing-based ST datasets are well suited for Spotiphy, mainly because of their whole genome coverage. Other approaches have enhanced their resolution but at the expense of gene coverage. For example, Slide-Seq v1/v2 achieves a resolution of approximately 10 µm but provides gene coverage of only 10^1^∼10^3^ ^13,14^; Stereo-seq offers sub-cellular resolution spot size (0.2 µm) with a gene coverage of 10^2^∼10^3^ ^16^. Data generated through these approaches does not stand to benefit greatly from the application of Spotiphy. At present, pairing gene-coverage-prioritizing approaches (e.g., Visium, DBiT-Seq) with Spotiphy may yield the best outcomes. Indeed, the data discussed in the Results section had a relatively high gene coverage of 10^3^∼10^4^ at single-cell resolution (Supplementary Fig. 19b).

Spotiphy’s performance is also contingent on the quality of the scRNA-seq reference for its generative model. The matched scRNA-seq datasets from the same samples would be ideal to avoid batch effects. However, acquiring such matched datasets in the real world is both technically challenging and financially cumbersome. To accommodate unmatched datasets, Spotiphy assigns variables (batch prior) to each gene to model the differences between ST and scRNA-seq profiles, effectively mitigating batch effects on deconvolution and decomposition. Spotiphy also presets four additional hyperparameters, including the quantile parameter, fold change threshold, non-zero count cell percentage, and number of marker genes to ensure scRNA-seq reference accuracy and robustness (Supplementary Fig. 1). In fact, the scRNA-seq reference used for mouse brains was primarily sourced from the Allen Institute, demonstrating that Spotiphy can produce reliable deconvolution results even when the reference originates from non-matched tissues. This flexibility significantly broadens Spotiphy’s potential applications, allowing for more uses in various research contexts without the need for matched reference datasets.

Spotiphy produces single-cell spatial whole transcriptomic profiles, but it is not the only solution. Emerging methods like iSpatial^77^ and Liger^78^ aim to generate the same results from image-based ST data, which could be worth exploring, but with some caveats. The major limit of image-based ST approaches the low-plexity of their pre-designed gene panels^9^. These methods use scRNA-seq data to infer the expression of nontargeted genes, a solution that is complicated by the different measurements that scRNA-seq and image-based ST use to quantify gene expression. For scRNA-seq, the raw count is a measure of the free mRNA molecules within an individual cell^79^, whereas the raw counts produced by image-based ST are a measure of the positive fluorescence signals within regions around the nuclei^80^. This fundamental discrepancy significantly diminishes the usefulness of scRNA-seq data as a reference and minimizing these "technique-derived batch effects" is challenging. Additional limitations make image-based ST data less suitable for inference. First, the fluorescence signal intensity is highly influenced by probe affinity, which varies across probe sets. Second, different excitation wavelengths will exhibit different signal intensities, complicating signal quantification. Third, image-based ST data tends to be sparser due to differences in the sensitivity of different probes. Altogether, these issues make it more difficult to accurately compare the expression levels between different genes.

Potential future directions for Spotiphy include the implementation of four improvements. First, a deep learning-based vision transformer model^39^ that correlates gene expression with image features would improve Spotiphy’s imputation function by taking advantage of extra information from histological images. This solution, which requires the use of massive training datasets, is contingent on the increased availability of sequencing-based ST data in the future^10^. It is also worth noting that for users studying cell-cell interactions at a fine scale, vision transformer modeling has the potential to address this issue. By leveraging the extensive learning process, the cell type information and iscRNA profile can be effectively correlated with the corresponding image features for each cell. Consequently, Spotiphy assigns a cell type and corresponding iscRNA profile to a random nucleus within each spot. For non-capture areas, Spotiphy assigns a cell type to each nucleus by randomly sampling based on the estimated cell type proportions. Although this random assignment is confined to a small region (50∼100 µm), it poses significant limitations in improving the accuracy of cell type assignment. A second exciting extension for Spotiphy involves reconstructing tissue’s 3D structure. Recent studies have shown that profiling sequential sections with spatial transcriptomics can reveal the real 3D organization of tissues^81,82^. Yet, it is costly and not feasible for broad use, where Spotiphy can step in. By leveraging the similarities in image features from adjacent sections, Spotiphy could infer the cellular proportions and expression profiles of neighboring sections, ultimately reconstructing the 3D structure. The image features would provide additional information to enhance the across-layer imputations^39^. STalign^83^ would be of great assistance for ensuring alignments between different sections. Another opportunity for improvement resides in Spotiphy’s inability to identify cell types that are present in the ST data but not in the scRNA-seq reference. We believe this shortcoming owes primarily to the algorithm’s inability to discern whether novel gene expression is indicative of an as-yet undefined cell type or simply a biological change in an existing cell type. In other words, the ease and accuracy of predictions could be improved if the gene coverage of sequencing allowed for a clearer distinction between cell subtypes and even substates. We therefore recommend including as many cell types as possible in the construction of the scRNA-seq reference so as to optimize the precision of Spotiphy’s results. Furthermore, with the continuous advancement of single-cell sequencing technologies, we aim to enhance Spotiphy by incorporating inputs from scMultiome (scRNA and scATAC)^84^ and Paired-Tag (scRNA and scChIP)^85^ profiles with spatial-ATAC-seq^86^ and spatial-CUT&Tag^87^, which have yielded spatially resolved chromatin modification and epigenome profiles. This expansion would enable Spotiphy to integrate and analyze multi-modal data, providing a more comprehensive and detailed understanding of spatially resolved cellular states and regulatory mechanisms. Lastly, the improvement in the resolution of ST platforms holds the potential to enhance existing methods that directly use ST data for principal component analysis and clustering analysis^88,89^. By leveraging the iscRNA data generated by Spotiphy as inputs, these methods will achieve more accurate and comprehensive output results at both gene and cellular levels. These outputs, including DEGs and neighborhood signatures, will provide more realistic guidance for future experimental validations.

---

**Extended Data Fig. 1.**
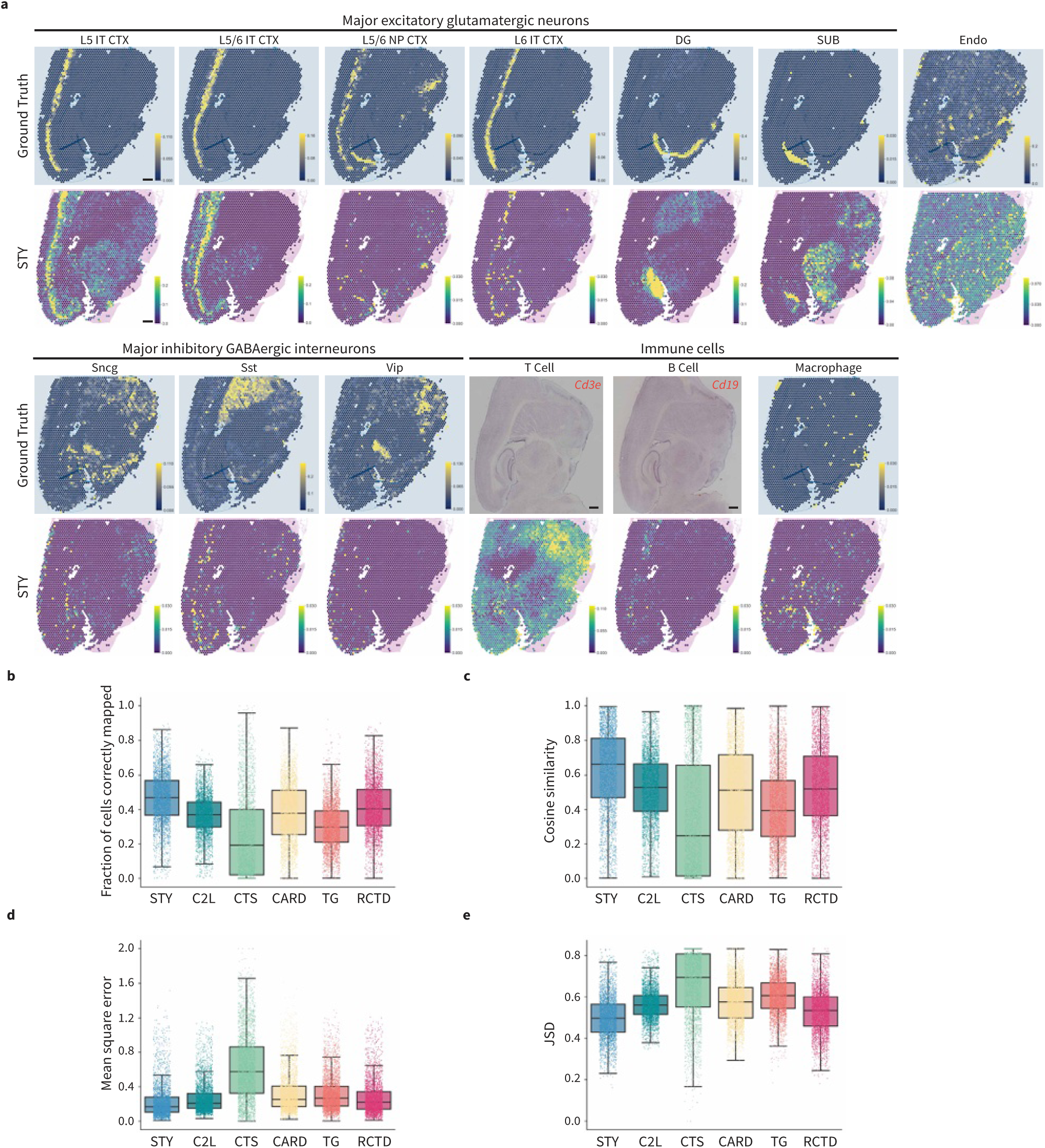
Spotiphy provides cell-type proportions that matches ground truth with high confidence. **Continued Fig. 2. a,** Heatmaps depicting the proportion of the rest cell types (as determined by Spotiphy and Xenium, the ground truth) across the histological image of AD sample. Scale bar: 500 μm. **b-e**, Box plots for the fraction of cells correctly mapped (**b**, higher is better), cosine similarity (**c**, higher is better), square error (**d**, lower is better), and JSD (**e**, Jensen–Shannon divergence, lower is better) of the cell-type proportions generated by each method for each transcriptomic spot.

**Extended Data Fig. 2.**
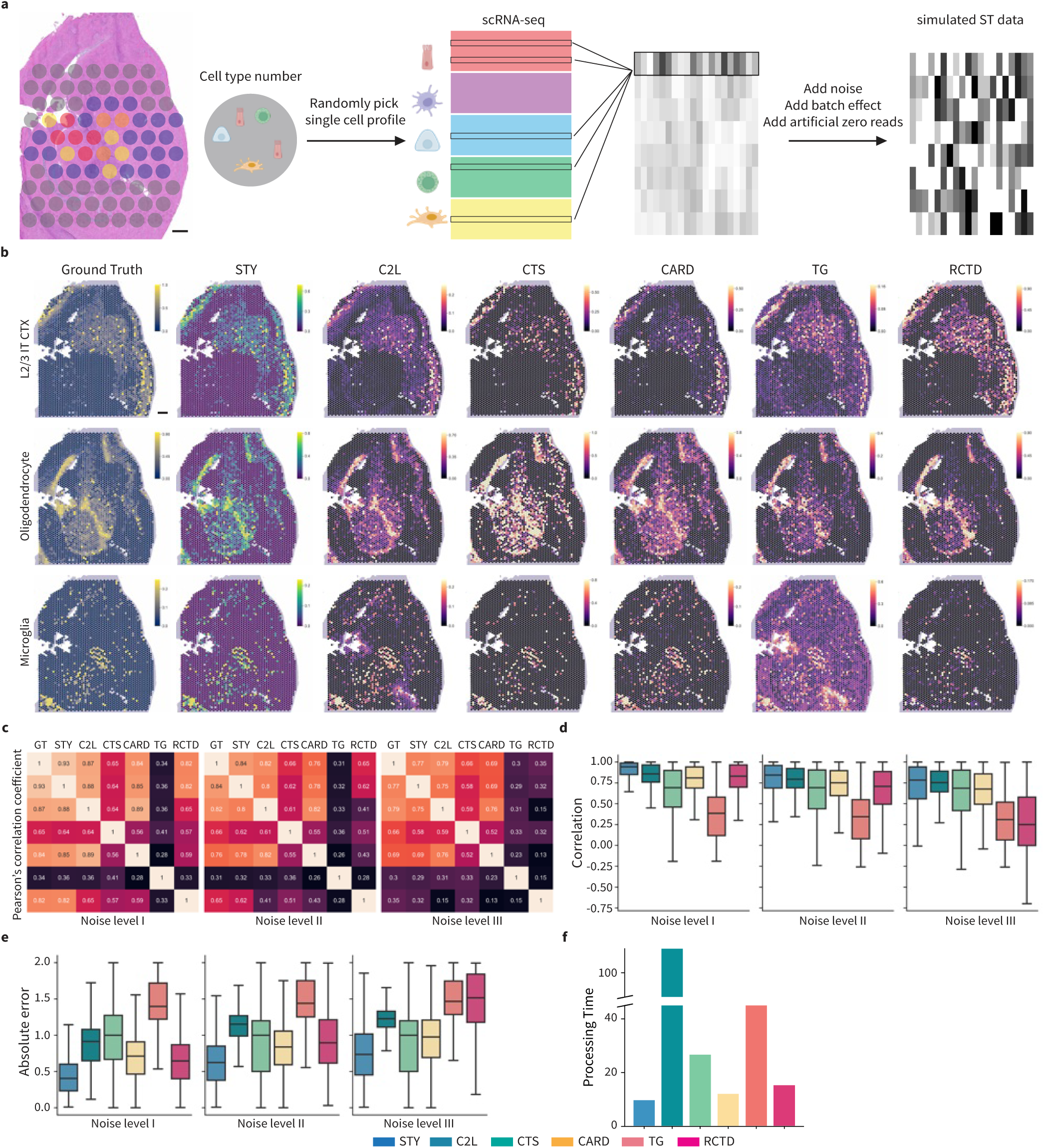
Simulated ST dataset further validates the superiority of Spotiphy. **a,** Workflow summary for simulated ST data generation based on deconvolution result of another mouse sample. Scale bars: 500 μm. **b**, Heatmaps depicting the proportion of three selected cell types (as determined by Spotiphy and Xenium, the ground truth) across the histological image of this mouse sample. Scale bars: 500 μm. **c**, Pearson correlation coefficient heatmap of cell-type proportions generated by Xenium, Spotiphy, and other methods selected for benchmarking. **d-e**, Box plots for the correlation (**d**, higher is better) and absolute error (**e**, lower is better) of the cell-type proportions for each transcriptomic spot generated by each method. **f**, Bar plot for processing time of all benchmarking methods.

**Extended Data Fig. 3.**
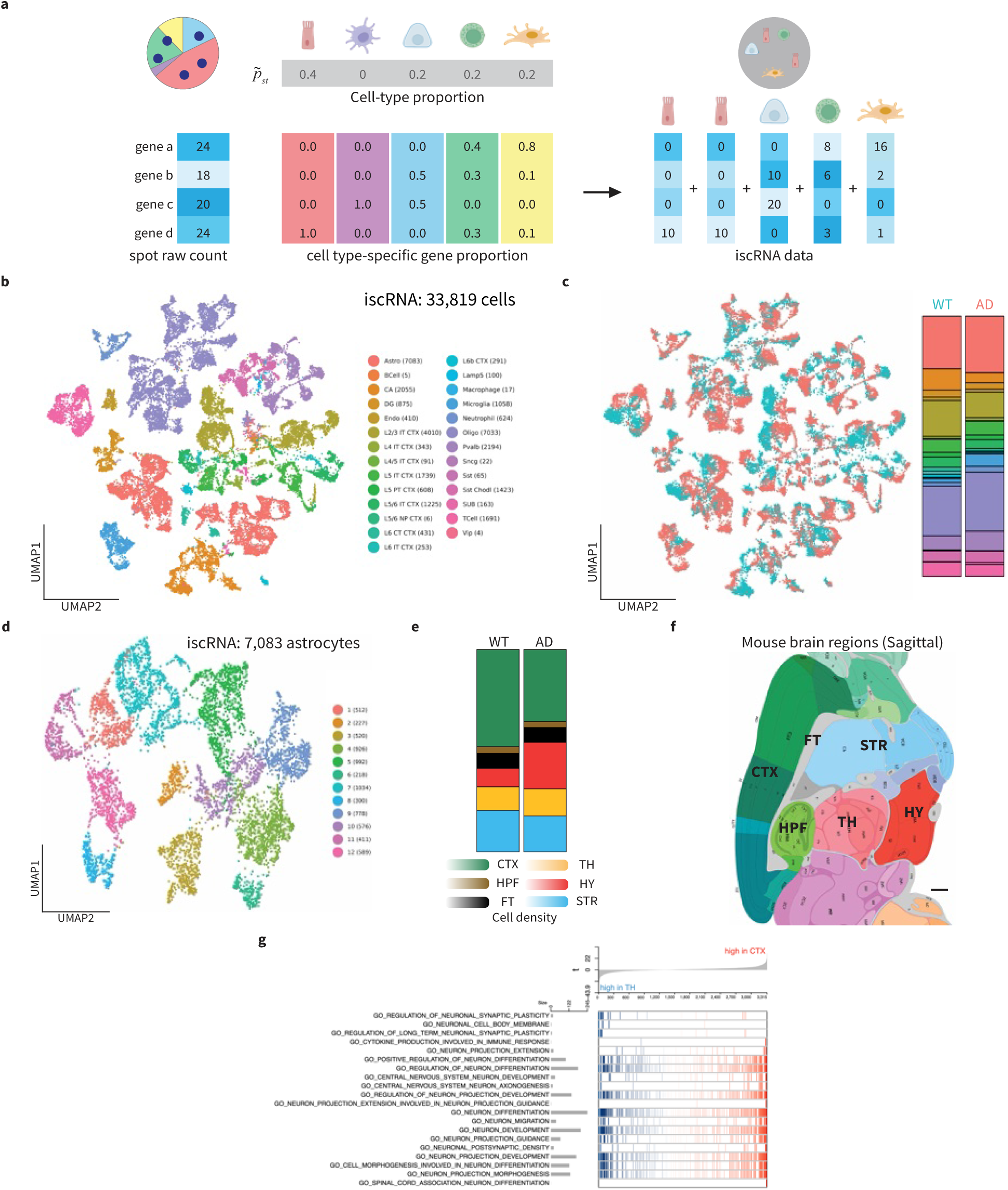
Decomposition outputs of Spotiphy and iscRNA data construction. **a,** Workflow summary for ST data decomposition and iscRNA data construction. **b-c**, UMAP projection of 33,819 cells from iscRNA data produced by applying Spotiphy to mouse brain Visium data. Clusters are labeled according to cell type (**b**) or sample-of-origin (**c**). **d,** UMAP projection of 7,083 astrocytes extracted from iscRNA data produced by applying Spotiphy to mouse brain Visium data. Clusters are labeled according to the original clusters. **e**, Bar plots of astrocyte subtype proportions annotated from Fig. 3a for both WT and AD mouse samples. **f**, Diagram for sagittal section of mouse brain labeled with different topographic regions. CTX: cerebral cortex; HPF: hippocampus; FT: fiber tracts; TH: thalamus; HY: hypothalamus; STR: striatum. Scale bars: 500 μm. **g**, Result of applying NetBID2 GSEA analysis to iscRNA data of astrocytes in CTX vs. TH.

**Extended Data Fig. 4.**
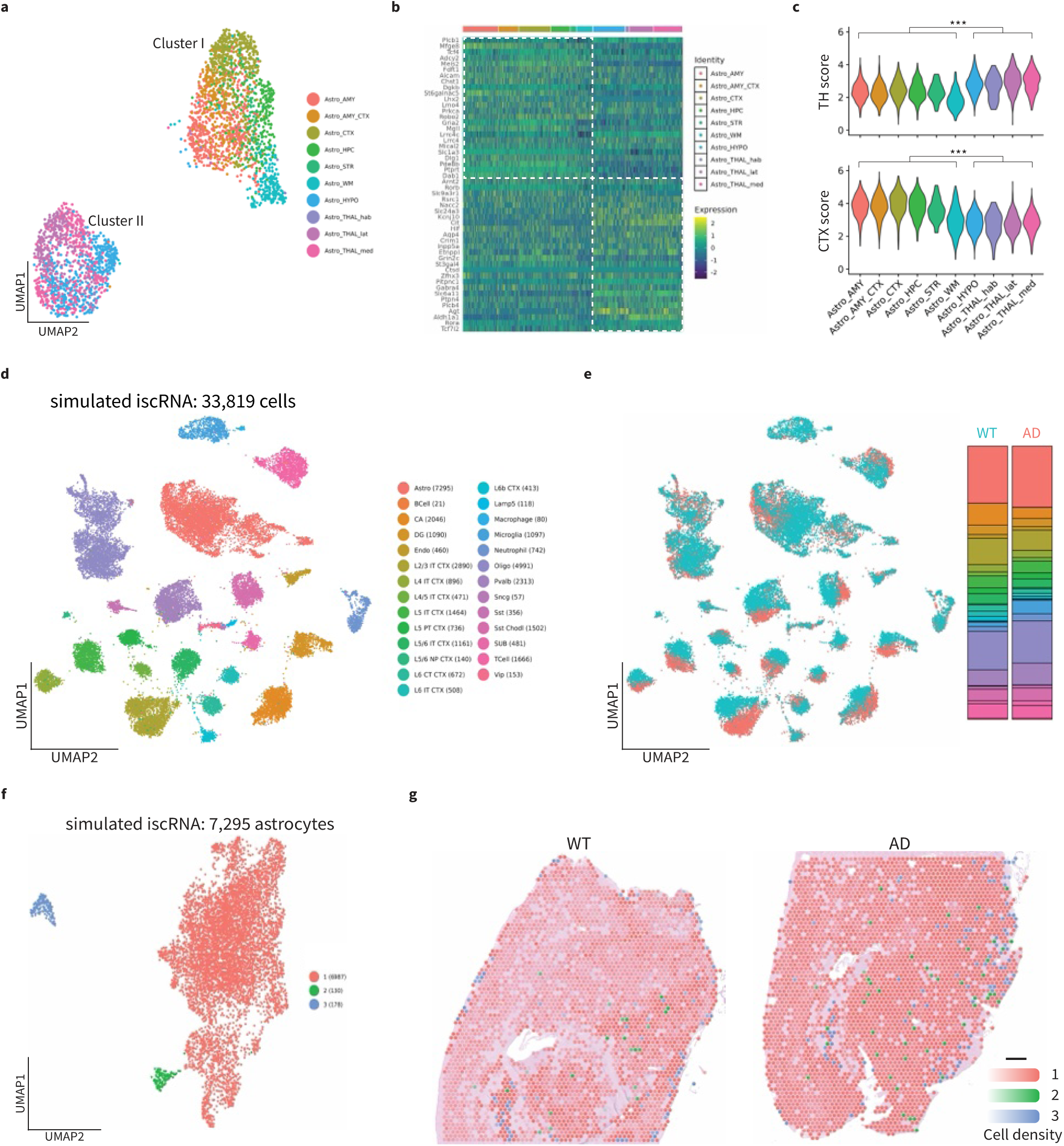
Prior research provides evidence validating astrocyte regional specification identified by iscRNA data generated by Spotiphy. **a-c**, Published astrocyte dataset^28^ confirmed astrocyte regional specification observed in iscRNA data. **a**, UMAP projection of published astrocyte dataset. **b**, Heatmap of expression of top DEGs identified from Fig. 3d among astrocytes in the published dataset. **c**, Violin plots of the CTX and TH scores for astrocytes in the published dataset. **d-g,** astrocytes from simulated iscRNA data lose regional specification. **d-e**, UMAP projection of 33,819 cells from simulated iscRNA data. Clusters are labeled according to cell type (**b**) or sample-of-origin (**c**). **f**, UMAP projection of 7,083 astrocytes extracted from simulated iscRNA data labeled with default clusters. **g**, Transcriptomic spots from the Visium data. Spots are color-coded according to their default clusters, annotated in **f**. The opacity of the spots is representative of cell density. Scale bars: 500 μm.

**Extended Data Fig. 5.**
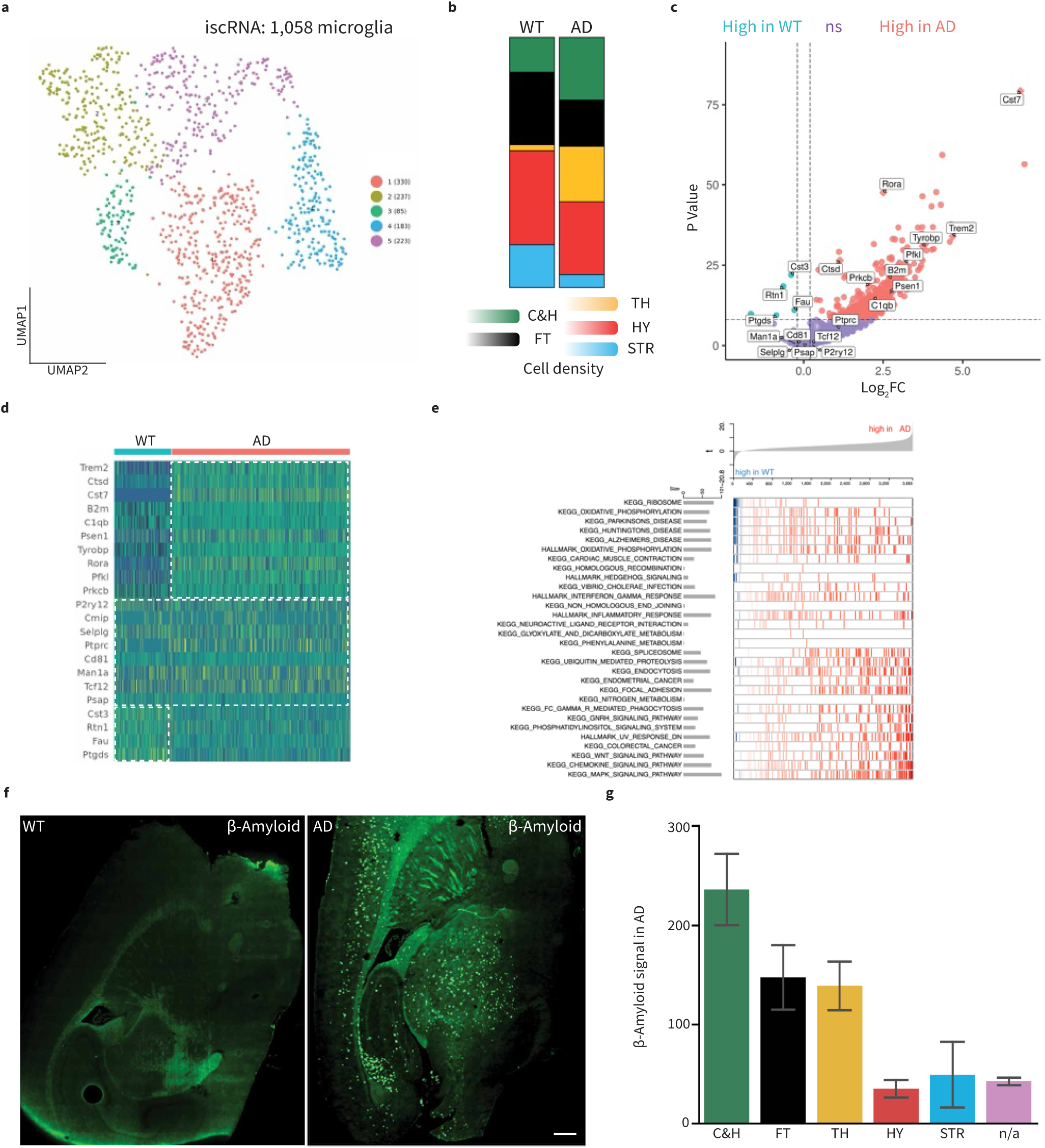
Unsupervised clustering analysis of iscRNA data identified DAM sub-populations. **a,** UMAP projection of 1,058 microglia extracted from iscRNA data produced by applying Spotiphy to mouse brain Visium data. Clusters are labeled with default clusters. **b**, Bar plots of microglia subtype proportions annotated from Fig. 4a for both WT and AD mouse samples. **c**, Volcano plot of differentially expressed genes (DEGs) among microglia in AD vs. WT. DEGs were determined using the iscRNA data. The genes highlighted are available in the CosMx panel and have been used to generate the signature scores in Fig. 4c**-e**. **d**, Heatmap of the expression of the top DEGs among microglia in WT and AD samples, respectively. **e**, Result of applying NetBID2 GSEA analysis to iscRNA data of microglia in AD vs. WT. **f**, beta-amyloid IHC staining of WT and AD samples. Scale bars: 500 μm. **g**, Quantification of beta-amyloid signals within different topographic regions of AD sample. The vertical bar represents the standard error.

**Extended Data Fig. 6.**
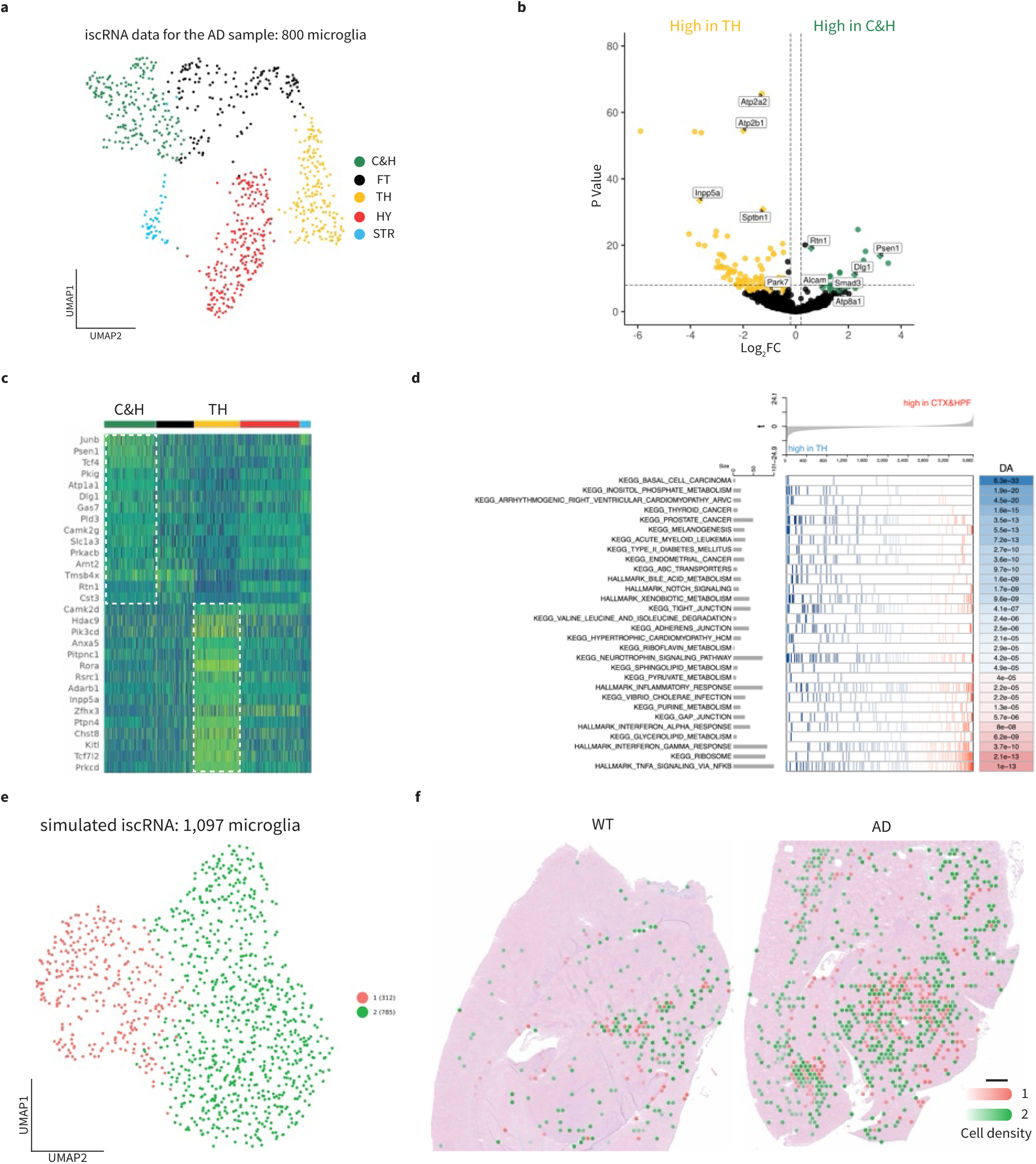
Spotiphy captures DAM regional specification in AD mouse brain tissue. **Continued Fig. 4. a,** UMAP projection of 800 microglia extracted from iscRNA data generated by Spotiphy to AD mouse brain Visium data. Clusters are labeled according to their corresponding topographic regions. **b,** Volcano plot of differentially expressed genes (DEGs) among microglia in C&H vs. TH. DEGs were determined using the iscRNA data. The genes highlighted are available in the CosMx panel and have been used to generate the signature scores in Fig. 4f**-h**. **c**, Heatmap of expression of top DEGs among microglia in AD sample. **d**, Result of applying NetBID2 GSEA analysis to iscRNA data of AD microglia in C&H vs. TH. **e**, UMAP projection of 1,093 microglia extracted from simulated iscRNA data labeled with default clusters. **f**, Transcriptomic spots from the Visium data. Spots are color-coded according to their default clusters, annotated in **e**. The opacity of the spots is representative of cell density. Scale bars: 500 μm.

**Extended Data Fig. 7.**
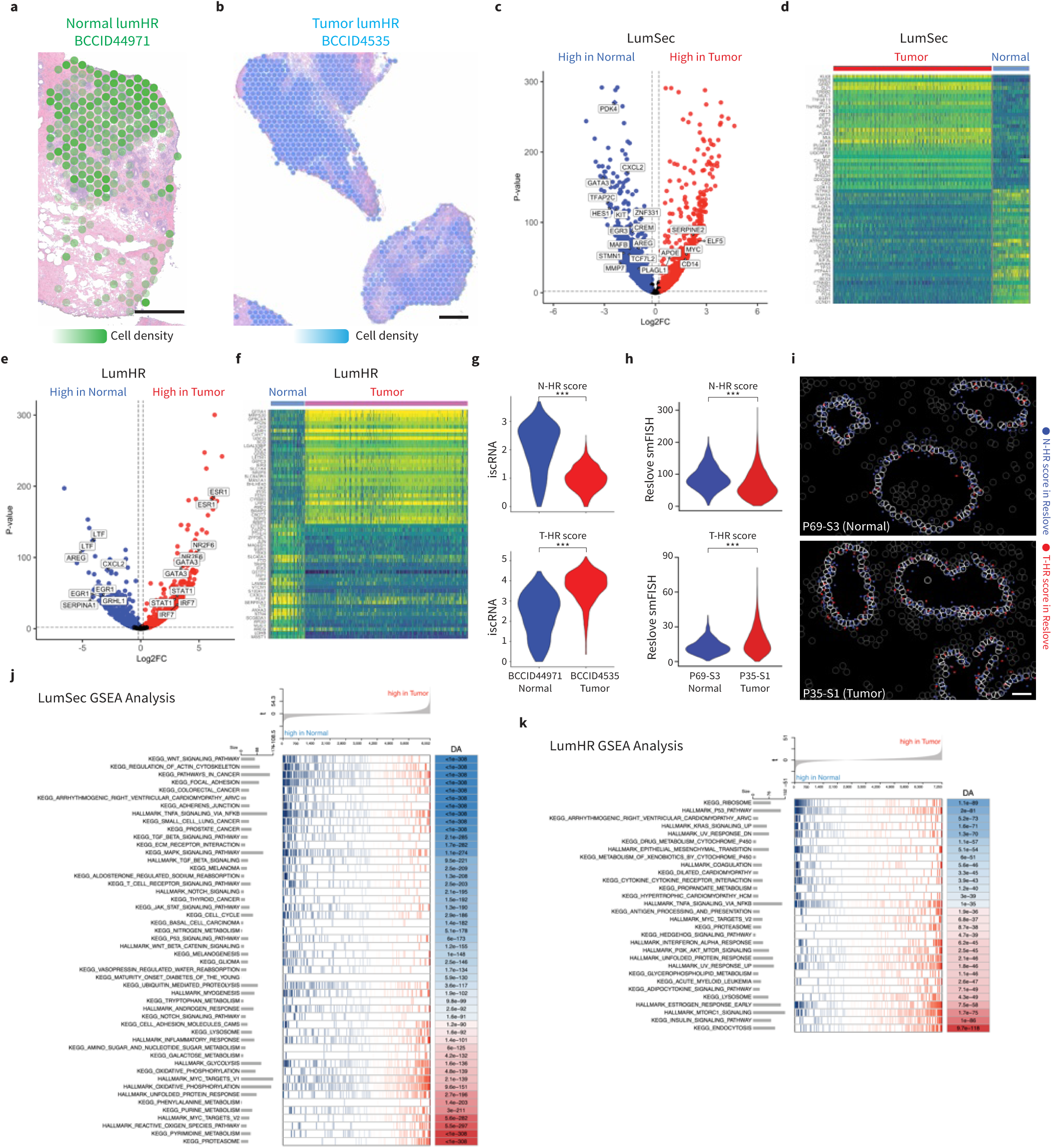
Spotiphy reveals tumor-microenvironment changes across spatial domains. **Continued Fig. 5. a-b**, Transcriptomic spots from the Visium data. Spots are color-coded according to their constituent LumHR cells’ sample-of-origin: BCCID4491 (**a**) and BCCID4535 (**b**) annotated in Fig. 5b. The opacity of the spots is representative of cell density. Scale bars: 500 μm. **c,** Volcano plot of differentially expressed genes (DEGs) among LumSec cells in BC1160920F vs. BCCID44971. DEGs were determined using the iscRNA data. The genes highlighted are available in the Resolve panel and have been used to generate the N-Sec and T-Sec scores in Fig. 5f**-h**. **c**, Heatmap of expression of top DEGs among LumSec cells. **e,** Volcano plot of DEGs among LumHR cells in BCCID4535 vs. BCCID44971. DEGs were determined using the iscRNA data. The genes highlighted are available in the Resolve panel and have been used to generate the N-HR and T-HR scores in **g-i**. **f**, Heatmap of expression of top DEGs among LumHR cells. **g,** Violin plots of the N-HR and T-HR scores for the LumHR cells identified in the iscRNA data of normal sample BCCID44971 and tumor sample BCCID4535. **h**, Violin plots of the N-HR and T-HR scores for the LumSec cells identified in the Resolve smFISH data of normal sample P69-S3 and tumor sample P35-S1. Two-sided t-tests are conducted. ***, p < 0.01. **i**, Visualization of signature genes in selected regions of the Resolve smFISH data. Blue and red represent the signature genes used to calculate the N-HR and T-HR scores, respectively. Cell boundaries are depicted with gray lines; LumSec cells are outlined in white. Scale bars: 100 μm. **j,** Result of applying NetBID2 GSEA analysis to iscRNA data of LumSec cells in BC1160920F vs. BCCID44971. **k,** Result of applying NetBID2 GSEA analysis to iscRNA data of LumHR cells in BCCID4535 vs. BCCID44971.

**Extended Data Fig. 8.**
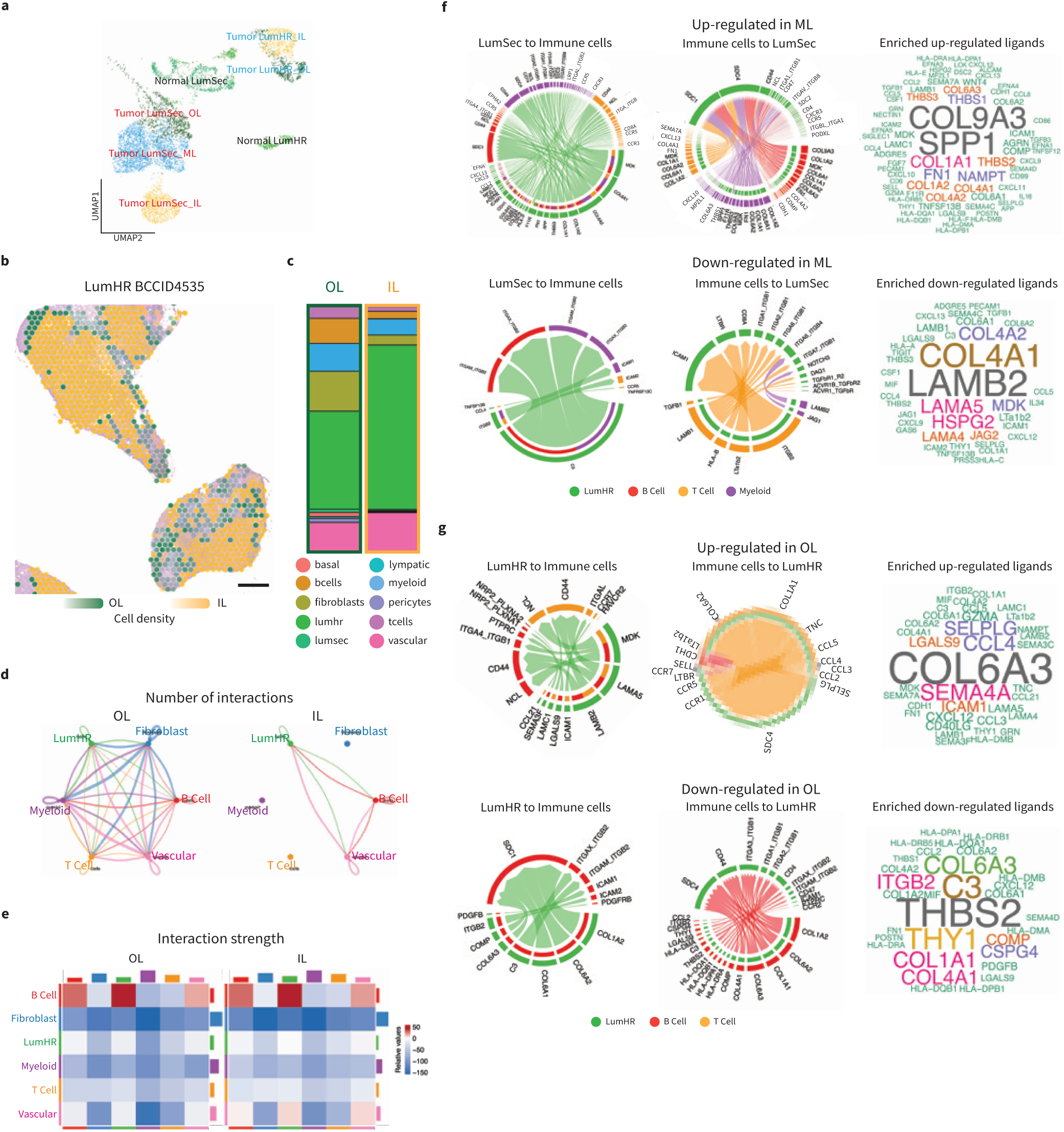
Downstream analysis of cell-cell communication patterns in human breast samples. **a,** UMAP projection of LumSec and LumHR cells from human breast tissue iscRNA data produced by applying Spotiphy to Visium data. Cells are color-coded according to their constituent LumSec cells’ and LumHR cells’ spatial domains. **b,** Transcriptomic spots from the Visium data. Spots are color-coded according to their constituent LumHR cells’ spatial domains. The opacity of the spots is representative of cell density. Scale bars: 500 μm. **c**, Cell-type proportions of the two spatial domains based on the iscRNA data shown in Fig. 5a. **d-e**, Results of applying CellChat to the iscRNA data of the cells in the three LumHR spatial domains. **d,** Number of interactions among cell types across three spatial domains. **e,** Interaction strength among cell types across three spatial domains. OL, outer layer; IL, inner layer. **f-g**, Detailed visualization of the up-regulated and down-regulated signaling ligand-receptor pairs using Chord diagrams of LumSec cells in ML domain (**f**) and LumHR cells in IL domain (**g**).

**Extended Data Fig. 9.**
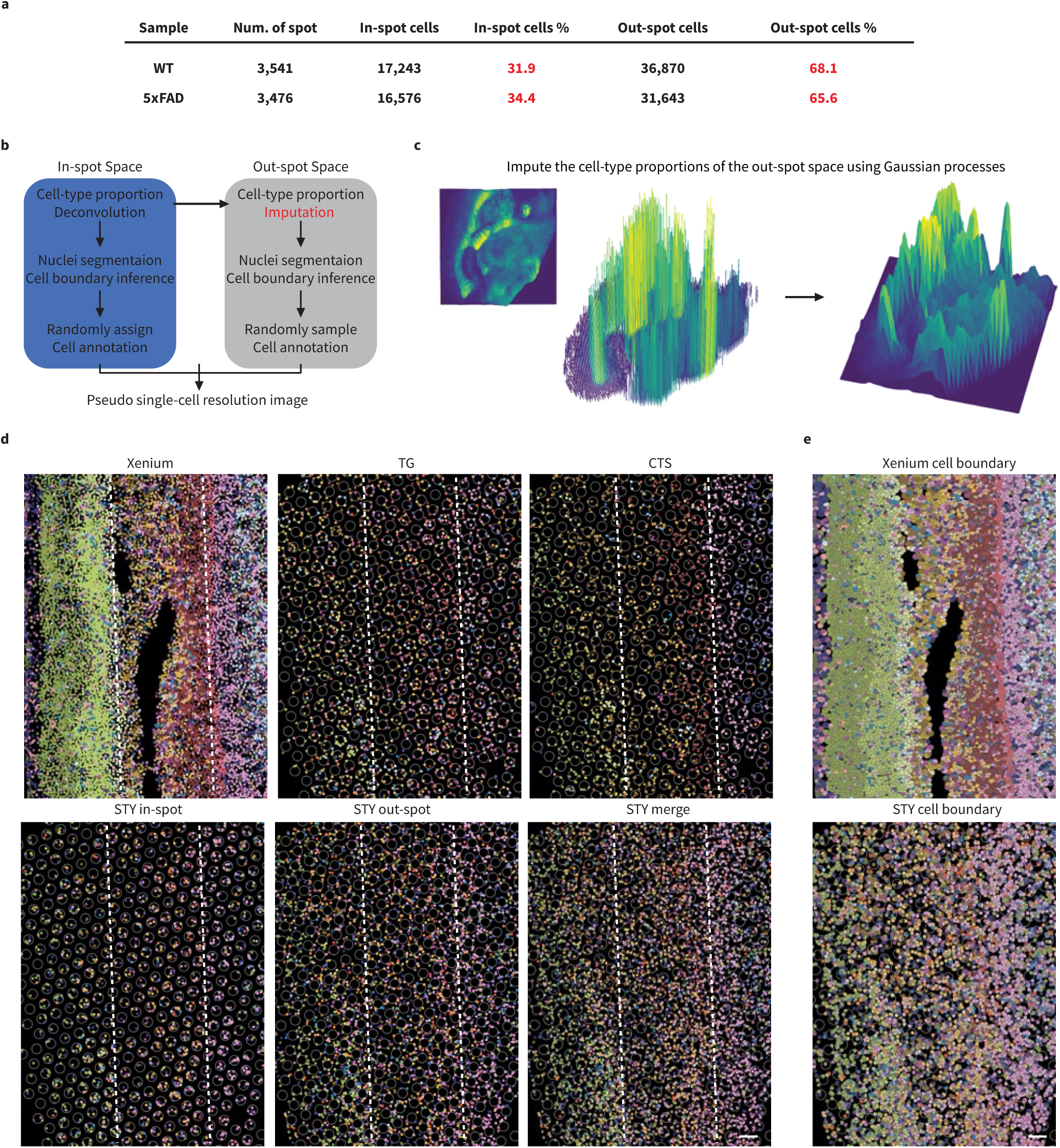
Spotiphy’s imputation to out-spot increases information density of ST data. **a,** Quantification of all detectable nuclei/cells including both in-spot and non-capture areas from WT and AD samples. **b**, The workflow of pseudo single-cell resolution image generation. **c**, Schematic overview of Gaussian processes for cell-type proportion imputation. **d-e,** Cell type annotation of individual cells in selected FOVs of cerebral cortex. Dots in **d** represent all detectable nuceli, color-coded for cell type. Grey circles represent the transcriptomic spots in the Visium data. Dashed white lines have been added to allow for easier comparison. **e,** Depicts whole cells, color-coded for cell type. The cell boundaries depicted for the Xenium (upper panel) and Spotiphy (lower panel) data were inferred through those methods’ respective pipelines. Color legend in Fig. 6. Scale bars in **d-e**: 100 μm.

**Extended Data Fig. 10.**
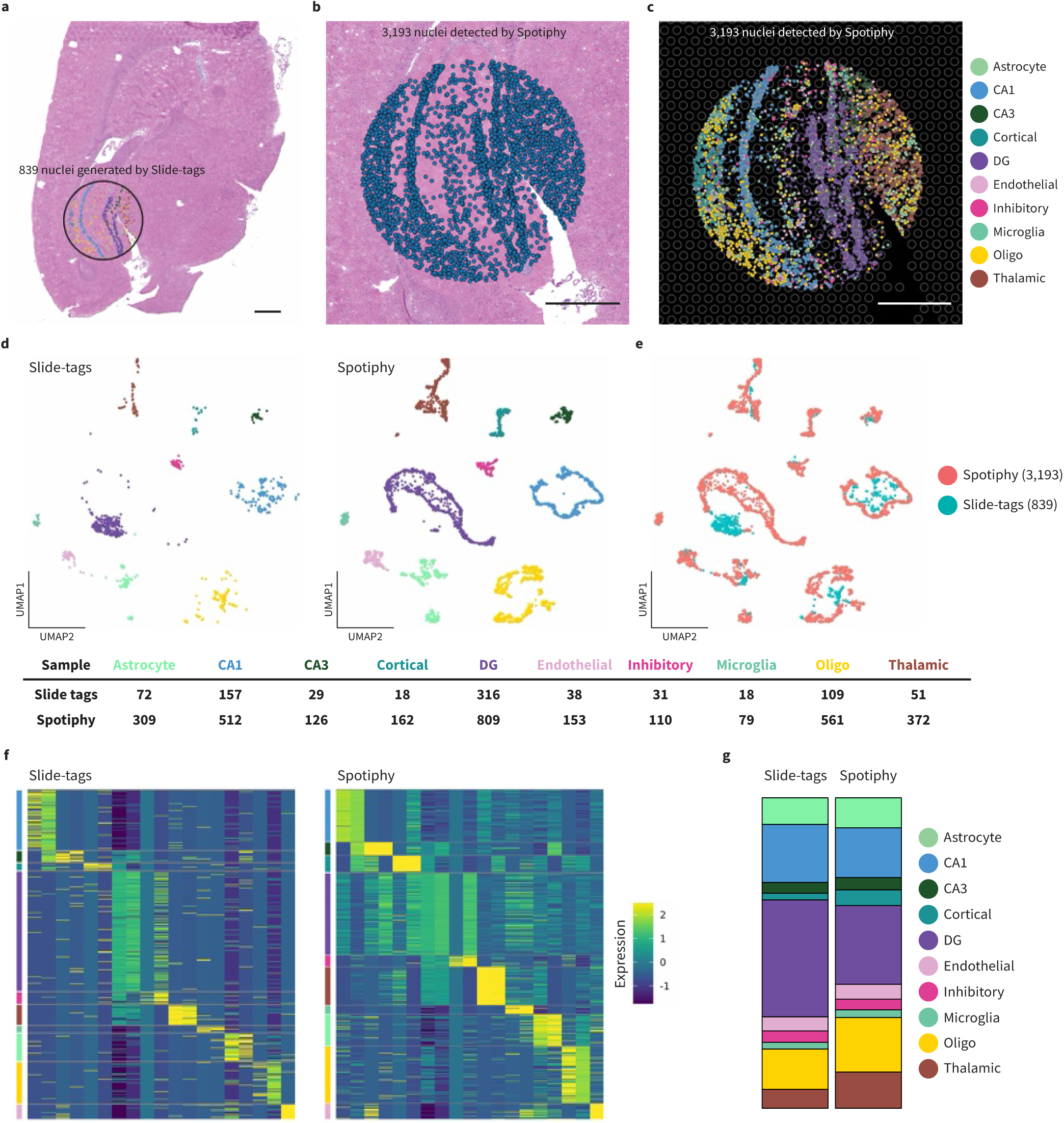
Spotiphy’s imputation to loss nuclei increases information density of Slide-tags data. **a,** Alignment of AD sample Visium H&E image and hippocampus plot from Slide-tags data of. **b-c**, Closer look at same hippocampus region of AD sample with 3193 nuclei detected by Spotiphy segmentation. **b**, nuclei are color-coded with blue to indicate their location on the H&E image. **c**, nuclei are color-coded for cell type annotated by Spotiphy. Grey circles represent transcriptomic spots from Visium data. Scale bars in **a-c**: 500 μm. **d-e**, UMAP projection of 4,032 cells including Slide-tags data and iscRNA data. Clusters are labeled according to their cell type (**d**) or data-of-origin (**e**). **f**, Heatmaps displaying the expression patterns of the marker gene sets from the original article in Slide-tags data and iscRNA data. **g**, Cell-type proportions of the Slide-tags data and iscRNA data.

## Data availability

The scRNA-seq datasets for mouse brains contain two parts: the original Allen Institute dataset is available at NeMO (https://assets.nemoarchive.org/dat-qg7n1b0), immune cell-enriched dataset generated in this study and final merged dataset used as reference are available at Zenodo data repository under record number 10520022. The matched ST datasets for mouse brains including Visium, Xenium, and CosMx are available at Zenodo data repository under record number 10520022. iscRNA datasets and simulated ST datasets used for evaluations are available at https://spotiphy.stjude.org. The snRNA-seq dataset for astrocyte was downloaded from the ArrayExpress under accession number E-MTAB-11115. The scRNA-seq dataset for human breast cancer was downloaded from the Gene Expression Omnibus under accession number GSE195665. The Visium datasets for human breast cancer were downloaded from the Zenodo data repository under record number 4739739.

## Code availability

Spotiphy version 1.0 was coded in Python and used to generate the results in this work. The source code for Spotiphy is freely available online at GitHub: https://github.com/jyyulab/Spotiphy. The documentation and tutorial with test inputs are available online at https://spotiphy.stjude.org.

## Author Information

These authors contributed equally: Jiyuan Yang, Ziqian Zheng

